# PP2A-dependent internalisation of GABA_B_ receptors in somatostatin interneurons regulates function and plasticity

**DOI:** 10.64898/2026.03.10.710829

**Authors:** Nithya Sethumadhavan, Max A Wilson, Anna Sumera, Desiree Loreth, Rita M Loureiro, Imre Vida, Akos Kulik, Sam A Booker

**Author notes:** Co-corresponding authors: Sam A Booker (Lead contact), Simons Initiative for the Developing Brain, Centre for Discovery Brain Sciences, University of Edinburgh, Hugh Robson Building, George Square, Edinburgh, EH8 9XD, UK, Akos Kulik, Institute of Physiology II, Faculty of Medicine, University of Freiburg, Hermann-Herder-Str. 7, Freiburg, 79104, Germany.

## Abstract

Cortical circuits rely on a precise balance of inhibitory and excitatory neurotransmission to encode information reliably and prevent pathology. Metabotropic GABA_B_ receptors (GABA_B_Rs) are key regulators of inhibitory signalling in mammalian neurons. In GABAergic interneurons (INs), GABA_B_R activation reduces inhibition overall, leading to disinhibitory mechanisms. In the hippocampus, somatostatin-expressing (SST) INs form a major subtype that provides feedback inhibition to the distal dendrites of principal cells (PCs) and other INs. Plasticity of SST INs is well established as a mechanism controlling hippocampal circuit function through both inhibitory and disinhibitory pathways and depends on metabotropic glutamate receptors (mGluRs) and GABA_B_Rs. However, whether activation of GABABRs induces metaplastic changes in SST INs, and how this influences circuit function and behaviour, remains unclear.

Here, we combined quantitative SDS-digested freeze-fracture replica immunoelectron microscopy, ex vivo electrophysiology, in vivo behavioural testing, and pharmacological manipulation of GABA_B_Rs. We show that receptor activation directly regulates SST IN plasticity via protein phosphatase 2A (PP2A)-dependent internalisation. GABA_B_R activation not only controls its own surface expression but also regulates membrane levels of mGluR1α and high-voltage-activated Ca_v_1.2 (L-type) Ca^2+^ channels. This GABA_B_R-dependent metaplasticity shifts circuit plasticity toward greater enhancement of long-range inputs to the CA1 region and disrupts contextual memory formation. These findings demonstrate that receptor-mediated surface dynamics in SST INs are critical for maintaining physiological neurotransmission and proper hippocampal microcircuit function.

## Introduction

The appropriate function of cortical microcircuits relies on dynamically balanced excitatory and inhibitory synaptic transmission, which maintains the neural code and contributes to memory formation^1–3^. How these circuits respond to changing activity level, or indeed sustained pharmacological activity, is crucial to understand how the brain functions at rest and how neuropathological conditions lead to altered behaviour and cognition.

Synaptic inhibition is produced, in large part, by a heterogenous population of local interneurons (INs). These INs release GABA from their presynaptic terminals, which activates both synaptic and extrasynaptic ionotropic GABA_A_ receptors (GABA_A_Rs) and metabotropic GABA_B_ receptors (GABA_B_Rs), on excitatory principal cells and INs alike ^4^. Such inhibition of INs reduces their activity, leading to reduced local GABA release, and thus a reduction of inhibitory control – known as disinhibition. Dynamic interactions between inhibition and disinhibition are critical to ensure the timing and strength of neuronal activity leading to complex behaviour. To date, most studied forms of disinhibition focus on direct synaptic GABA_A_R-mediated effects, which can regulate circuit level plasticity in hippocampus ^5^ and neocortex ^6^. While the direct effect of GABA_B_R signalling on INs has been well described by ourselves and others ^7–14^, how their activation shapes the long-term excitability of inhibitory cells and influence circuit activity remains less well understood.

In CA1 of the hippocampus, a major type of IN are those which express the neuropeptide somatostatin (SST)^15^ which have their somatodendritic domain within *stratum oriens/alveus* (str. O/A) receiving inputs from local pyramidal cells (PCs)^10,16,17^, with axons ramifying in *str. lacunosum-moleculare* (*Str. L-M*). Thus, SST INs provide powerful feedback inhibition into the CA1 circuit^18^. Therefore, the activation of SST INs leads to complex circuit functions. First, they directly inhibit distal dendrites of CA1 PCs co-terminating with entorhinal cortex inputs to strongly inhibit spatial information^19–21^. Second, SST INs disinhibit proximal dendritic domains and thus gate contextual inputs from CA3^22^. As such, long-term potentiation of SST INs leads to a functional re-arrangement of the hippocampal information transfer, favouring CA3 inputs ^22,23^ and are potentially able to alter the circuit dynamics of the hippocampus over behaviourally relevant time-scales ^24^. We have shown that GABA_B_Rs on SST INs acutely inhibit the induction of long-term potentiation (LTP)^11^ and their local circuit integration ^10^, but do not directly hyperpolarise SST INs. Given the high affinity of GABA for GABA_B_Rs and their plethora of accessory proteins, they undergo rapid desensitization ^8,25–29^ and agonist dependent internalisation ^30,31^. Not only this, but GABA_B_Rs are known to interact with metabotropic glutamate receptors (mGluRs), preferentially with type 1α receptors (mGluR1α), in other GABAergic cells in the cerebellum ^32,33^. Considering the high density of both mGluR1α, and GABA_B_Rs on SST INs ^34,35^ it is plausible that receptor cross-talk also exists in these cells. Whether GABA_B_R signalling is susceptible to rapid internalisation in SST INs, and how this affects cell function and the local circuit remains unknown.

We hypothesised that GABA_B_R activation leads to internalisation of itself, favouring greater plasticity in SST INs. To determine this, we performed quantitative SDS-digested freeze-fracture replica immunogold labelling (SDS-FRL) electron microscopy, *ex vivo* electrophysiological recordings from CA1 of the mouse hippocampus following prolonged application of the selective GABA_B_R agonist baclofen, and behavioural characterisation. We found, in opposition to our hypothesis, that after sustained GABA_B_R activation, synaptic plasticity of SST INs was reduced, independent of direct receptor activation. This reduction in plasticity was due to concomitant protein phosphatase 2A (PP2A)-mediated internalisation of GABA_B_Rs, Ca_v_1.2 (L-type) voltage-gated Ca^2+^ channels (VGCCs) and mGluR1α – both of which are required for LTP of SST INs. Finally, we show that baclofen administration leads to circuit level strengthening of long-range inputs to CA1, which are dependent on SST IN plasticity, and an inability to form contextual memories.

## Results

In this study we tested the hypothesis that sustained baclofen application leads to internalisation of GABA_B_Rs, which favours enhanced plasticity of SST INs following receptor activation. For this, we performed detailed quantitative immunoelectron microscopy, electrophysiology, and behaviour in adult mice.

### Native GABA_B_Rs, mGluR1α and L-type calcium channels are down-regulated in SST IN membranes following baclofen pre-application, which requires PP2A

We have previously shown that mouse SST IN dendrites possess a high density of GABA_B1_, Ca_V_1.2, and mGluR1α^11^. The latter two proteins are required for associative theta-burst stimulus (aTBS) LTP of SST INs ^36,37^, and is inhibited by GABA_B_Rs ^11^. To determine whether pre-application of the GABA_B_R agonist baclofen (20 μM) altered membrane localization of GABA_B_Rs as compared to mGluR1α and Ca_V_1.2, we performed SDS-FRL immunoelectron microscopy analysis from acute mouse hippocampal slices pre-treated with baclofen (n=5 mice), compared slices treated with vehicle (n=5 mice) or the GABA_B_R antagonist CGP-55,845 (5 μM, n=3 mice; Figure 1A). All SDS-FRL data are shown as animal average values, with statistics shown based on linear mixed-effects modelling to account for inter-animal variability. Data from each dendritic profile within animal is shown in Figure S1.

**Figure 1.**
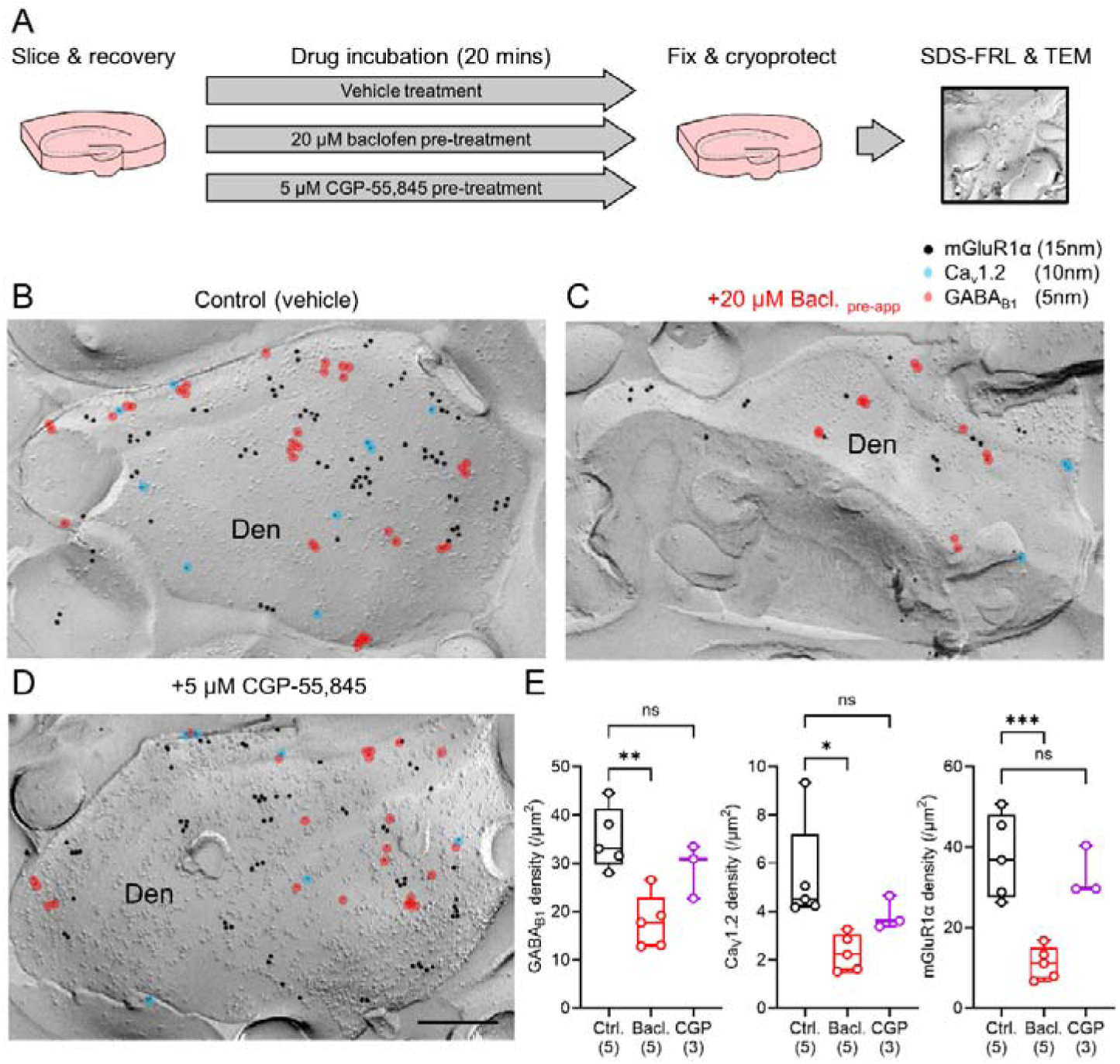
Sustained GABA_B_R activation leads to reduction of membrane associated GABA_B1,_ Ca_V_1.2, and mGluR1α proteins from putative SST-INs. **A** Schematic of the experimental overview, indicating treatment and experimental sample times. **B** Example electron micrographs of SDS-FRL for mGluR1α (15 nm immunoparticles), Ca_V_1.2 (10 nm, blue overlay) and GABA_B1_ (5 nm, red overlay) on a putative SST IN dendrite in *str. O/A* of CA1 in a vehicle slice. The same labelling but performed in brain slices following 20 minutes baclofen pre-application (20 μM, **C**) or CGP-55,845 (5 μM, **D**). **E** Quantification of labelling density for GABA_B1_ (left), Ca_v_1.2 (middle) and mGluR1α (right), in slices treated with vehicle (Ctrl., black, 5 mice), baclofen pre-application (Bacl., red, 5 mice), or CGP-55,845 (CGP, magenta, 3 mice). All data are shown as boxplots depicting the 25-75% range with median, maximum range and data from individual mice overlaid as open circles. Statistics shown as: ns – p>0.05, * - p<0.05, ** - p<0.01, *** - p<0.001; all from post hoc Tukey tests following LMM analysis. Scale bar represents 200 nm.

Under control conditions (Figure 1B), putative SST IN dendrites were identified based on high levels of surface labelling for mGluR1α ^34^, with an average density of 37.6 ± 4.7 particles/μm^2^ (Figure1B, E). Consistent with our previous data in rats ^11^, mGluR1α-positive dendrites displayed high-density GABA_B1_ subunit labelling of 35.1 ± 2.9 particles/μm^2^ and moderate expression of the L-type channel subunit Ca_V_1.2 at 5.46 ± 0.98 particles/μm^2^ (Figure 1B, E). Compared to control slices, baclofen pre-application led to a 49% reduction in density to 17.9 ± 2.5 particles/μm^2^ for GABA_B1_ (t_(5,5)_=5.07, *p*=0.004, Tukey test; Figure 1C, E). We observed a 58% reduction in Ca_V_1.2 to 2.3 ± 0.3 particles/μm^2^ (t_(5,5)_=5.36, *p*=0.0009, Tukey test) and a 70% reduction of mGluR1α labelling 11.2 ± 1.8 particles/μm^2^ (t_(5,5)_=6.36, *p*=0.0004, Tukey test) (Figure 1C, E). Application of the GABA_B_R antagonist CGP-55,845 did not decrease density of GABA_B1_ (t_(5,3)_=1.97, *p*=0.18, Tukey test), Ca_V_1.2 (t_(5,3)_=2.01, *p*=0.17, Tukey test) or mGluR1α (t_(5,3)_=0.69, *p*=0.78, Tukey test) relative to control levels (Figure 1D, E). This effect on effector channels appeared confined to SST INs, as measurement of CA1 PC dendrites in the same replicas revealed that while baclofen pre-application reduced GABA_B1_ density by 49.9% (F_(5,5)_=45.22, p=4.57×10^−10^, LMM, Supplementary Figure 2) Ca_V_1.2 was unchanged (F_(3,3)_=0.009, p=0.93, LMM; Supplementary Figure 2). mGluR1α was not tested as this protein is not expressed abundantly on CA1 PCs. Together, these data reveal that GABA_B_R activation leads to its own reduction from the membrane of SST INs, but also leads to down-regulation of Ca_V_1.2 and mGluR1α.

### Baclofen pre-application leads to reduced Cav1.2 and group 1 mGluR-mediated currents in SST INs

To determine whether baclofen pre-application regulates the function of Ca_v_1.2 and mGluR1α in SST INs, we next performed whole-cell patch clamp recordings from identified cells using targeted pharmacology to interrogate receptor internalisation (Figure 2A), as GABA_B_R internalisation has been shown to be dependent on increased activity of PP2A^26,31,38^.

**Figure 2:**
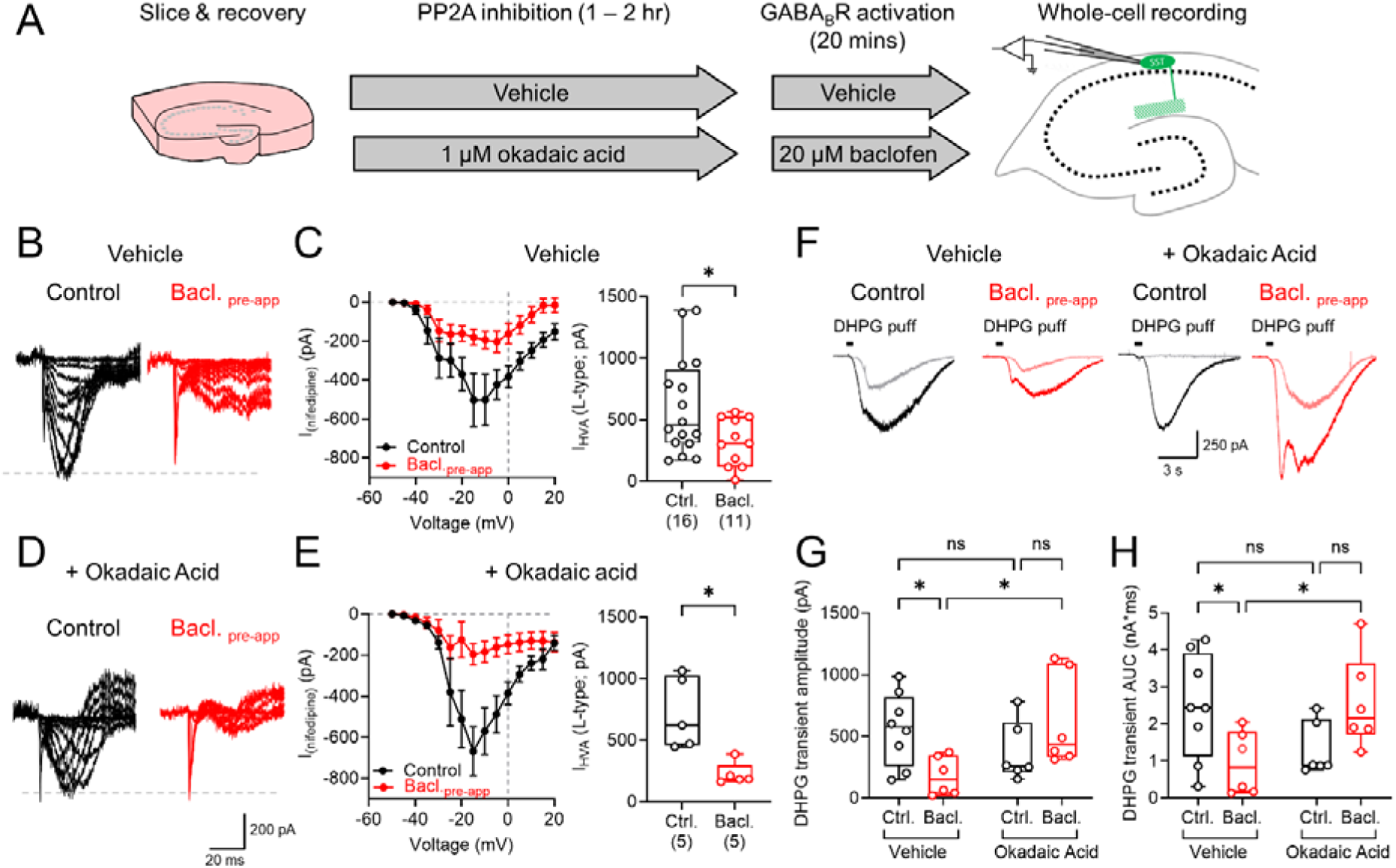
Ca_V_1.2 and group 1 mGluR-mediated currents are reduced following baclofen pre-application. **A** Schematic of recording conditions, highlighting treatment periods and recording times. **B** Example nifedipine-sensitive VGCC current responses recorded in SST INs following depolarising stimulation from −60 mV. Dashed line indicates control response amplitude for comparison. **C** Quantification of voltage-dependency of nifedipine-sensitive VGCC currents (left) and peak nifedipine-sensitive currents (right). **D** Example nifedipine-subtracted isolated VGCC current responses in SST INs pre-application with 1 μM okadaic acid. **E** Quantification of voltage-dependency of nifedipine-sensitive VGCC currents (left) and peak nifedipine-sensitive currents (right) in okadaic acid pre-application SST INs. **F** Example voltage responses following 50 μM S-DHPG puff application to SST INs, recorded under control conditions (black) or following baclofen pre-application (red), following either vehicle or 1 μM okadaic acid pre-application. **G** Quantification of absolute current amplitudes following S-DHPG puff. **H** Quantification of the area-under-curve (AUC) for S-DHPG puff induced transients. Data are shown as either mean ± SEM (current-voltage plots) or box-plots showing 25-75% box with median, with maximum range shown. Individual cell data is shown overlaid. Statistics shown as: ns – p>0.05, * - p<0.05; from 2-way ANOVA and Mann Whitney non-parametric tests

As Ca_v_1.2 containing VGCCs are a major target of GABA_B_R mediated inhibition in SST INs^11^, we first performed whole-cell recordings using a Cs-gluconate based internal solution containing QX-314 (5 mM) to block voltage activated K^+^ and Na^+^ channels, respectively, in the presence of CNQX (10 μM), DL-AP5 (50 μM) and gabazine (10 μM) to block ionotropic receptor signalling. High-voltage activated Ca^2+^ responses were measured from −60 mV voltage-clamp, in which 5 mV depolarising current steps were applied, followed by bath application of the selective L-type VGCC blocker nifedipine (10 μM; Figure 2B). The average subtracted L-type current measured across a range of depolarising stimuli revealed large amplitude currents in SST INs recorded under control conditions, which were substantially reduced, but with a similar voltage-dependency in slices pre-application with baclofen. We found that baclofen pre-application reduced the peak L-type current measured by 37% (U_(16,11)_= 50_;_ P=0.032, Mann-Whitney test, Figure 2C). We performed the same recordings, but in slices that had been incubated with the PP2A inhibitor okadaic acid (OA, 1 μM; Figure 2D). Blocking PP2A activity had minimal effect on current-voltage response or peak L-type currents following baclofen pre-application (U_(5,5)_= 0_;_ P=0.008, Mann-Whitney; Figure 2E).

For mGluR1 signalling, we recorded SST INs in whole-cell configuration then puff applied the selective, high-affinity group I mGluR agonist S-DHPG (50 μM, 200 ms, 10 PSI) within 100-200 μm of the cell body. S-DHPG puffs resulted in large and slow depolarising currents in SST INs, which were partially sensitive to the mGluR1α selective antagonist LY367,385 (100 μM). Measurement of the same S-DHPG induced currents in slices that were pre-application with baclofen possessed typically smaller currents (Figure 2F). Quantification of peak S-DHPG currents revealed that baclofen pre-application reduced the amplitude (F(1,22)=7.99, P=0.01, 2-way ANOVA [interaction], Figure 2G) and the charge transfer (F(1,22)=9.59, P=0.005, 2-way ANOVA [interaction], Figure 2H), effects which were not present when slices were incubated with OA prior to baclofen pre-application.

### Baclofen pre-application dependent reductions in Ca_v_1.2 and mGluR1α are mediated by PP2A

To confirm surface down-regulation of GABA_B_Rs was indeed dependent on PP2A in SST INs, we utilised a mouse line expressing a Channelrhodopsin2 fused to yellow fluorescence protein (ChR2/YFP) under the SST promotor to allow selective SDS-FRL from identified SST IN dendrites ^10^. Initially, we prepared slices from WT mice treated with the PP2A inhibitor OA. However, in these slices we observed minimal to no labelling for mGluR1α, GABA_B1_ or Ca_V_1.2 labelling in putative SST IN and CA1 PC dendritic profiles (Figure S3). This effect was observed across >5 independent experiments.

As such, we next used the the small molecule PP2A inhibitor fostriecin (50 nM) which slices were incubated with for 1 hour prior to baclofen pre-application (Figure 3A). In *str. O/A* from the ChR2/YFP mice, we observed strong immunolabeling for YFP (33.7 ± 1.07 particles/μm^2^, Figure 3B). In vehicle slices, we observed a 45% reduction of GABA_B1_ labelling following baclofen pre-treatment (16.7± 1.02 particles/μm^2^, t_(3,2)_=3.45, *p*=0.037, Kenward-Roger test, Figure 3B, D), consistent with observations from mGluR1α labelled dendrites (Figure 1). Incubation with fostriecin prevented the baclofen pre-treatement mediated reduction in GABA_B1_ surface density (28.3 ± 3.2 particles/μm^2^ t_(3,2)_=0.88, *p*=0.44, Kenward-Roger test, Figure 3C, D), confirming that PP2A inhibition prevents GABA_B_R internalisation.

**Figure 3:**
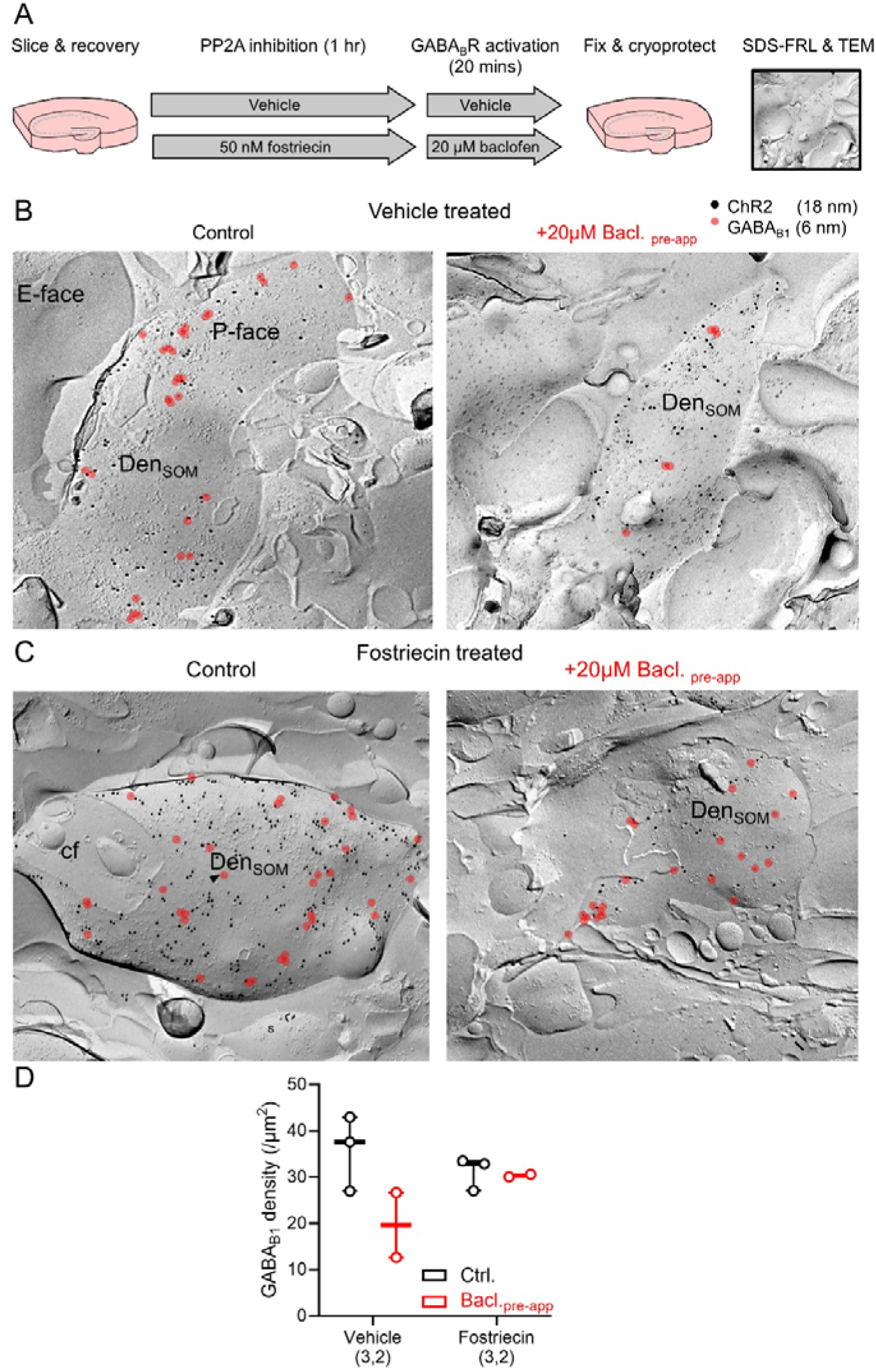
Inhibition of PP2A prevents internalisation of GABA_B_Rs in SST INs following their activation. **A** Schematic of experimental conditions, highlighting treatment periods and fixation times **B** Example electron micrographs of SDS-FRL for SST-cre dependent expression of ChR2/YFP (18 nm immunoparticles) and GABA_B1_ (6 nm, red overlay) in *str. O/A* of CA1 of slices with vehicle or baclofen pre-application (20 μM). **C** The same labelling but performed in slices incubated with 50 nM fostriecin for 1 hour prior to baclofen pre-application. **D** Quantification of GABA_B1_ density in vehicle slices (Ctrl., black, 3 mice) or those pre-application with baclofen (Bacl., red, 2 mice) with and without fostriecin incubation. All data are shown as boxplots depicting the 25-75% range with median, maximum range shown, and average density calculated from individual mice results overlaid as open circles. Scale bar represents 200 nm.

To determine whether GABA_B_R-dependent internalisation of L-type calcium channels also requires PP2A-dependent mechanisms, we performed the same experiments but labelling for Ca_V_1.2 (Figure 4A). In replicas from vehicle treated slices we observed Ca_V_1.2 labelling of 3.7± 0.3 particles/μm^2^, which was 46% reducted in baclofen pre-application slices (1.8 ± 0.3 particles/μm^2^ t_(3,2)_=3.32, *p*=0.0497, Kenward-Roger test, Figure 4B, D), consistent with with mGluR1α labelling experiments (Figure 1). Fostriecin pre-incubation (Figure 3C) prevented this reduction in Ca_V_1.2 labelling following baclofen treatment (3.5 ± 0.4 particles/μm^2^ t_(3,2)_=1.06, *p*=0.326, Kenward-Roger test, Figure 3C, D), confirming that PP2A inhibition prevents Ca_v_1.2 channel internalisation.

**Figure 4:**
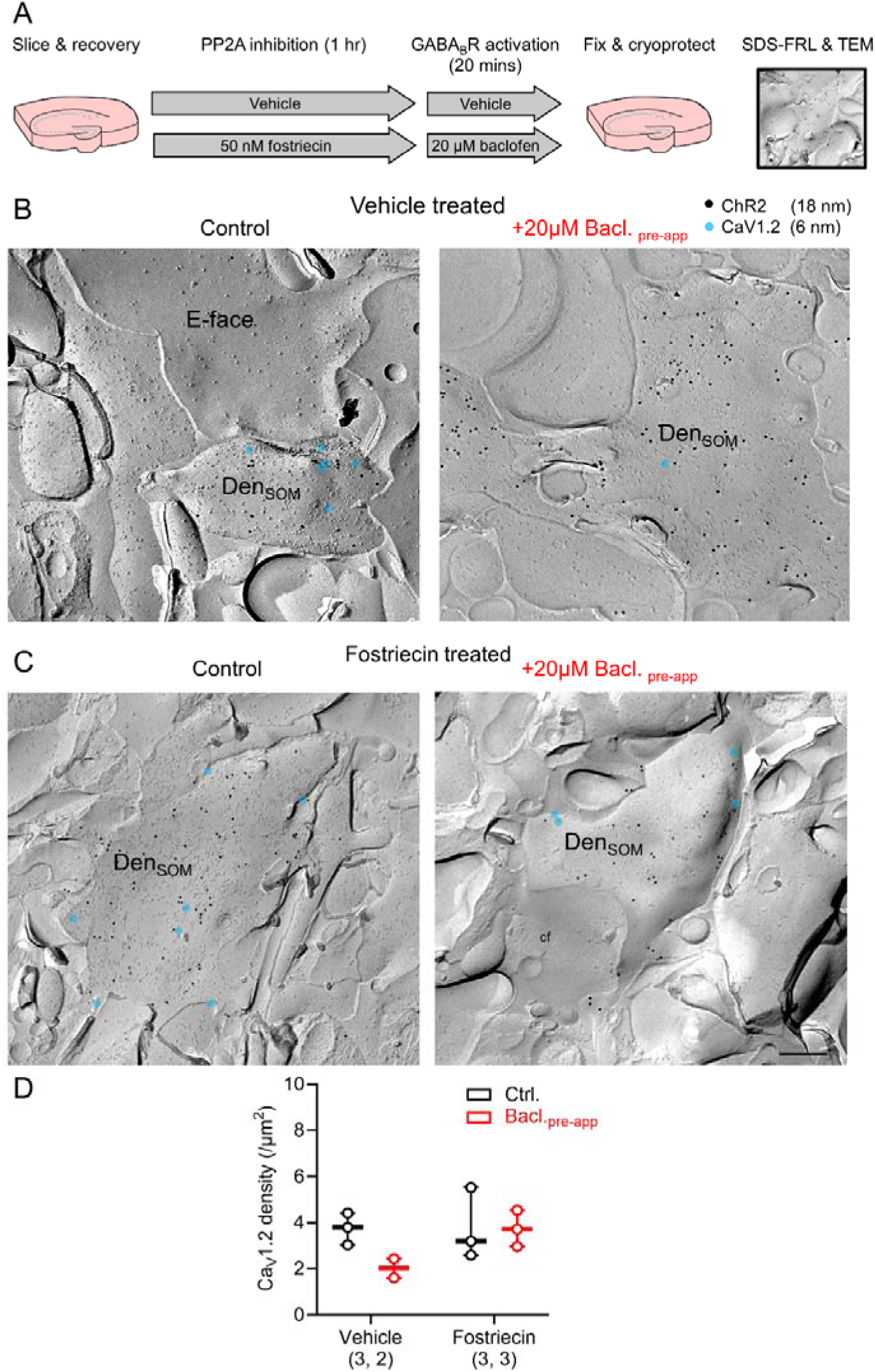
Inhibition of PP2A prevents internalisation of Ca_V_1.2 in SST INs following GABA_B_R activation. **A** Schematic of experimental conditions, highlighting treatment periods and fixation times. **B** Example electron micrographs of SDS-FRL for SST-cre dependent expression of ChR2/YFP (18 nm immunoparticles) and Ca_V_1.2 (6 nm, blue overlay) in *str. O/A* of CA1 of slices with vehicle or baclofen (20 μM). **C** The same labelling but performed in slices pre-application with 50 nM fostriecin for 1 hour prior to baclofen application. **D** Quantification of Ca_V_1.2 density in slices treated with vehicle (Ctrl., black, 3 mice) or baclofen (Bacl., red, 2 mice) with and without fostriecin pre-application. All data are shown as boxplots depicting the 25-75% range with median, maximum range shown, and average density calculated from individual mice results overlaid-as open circles. Scale bar represents 200 nm.

These data confirm that baclofen induced GABA_B_R internalisation requires PP2A in SST INs in *str. O/*A, and that inhibiting this internalisation also prevents loss of Ca_V_1.2 from their dendritic membranes.

### Sustained GABA_B_R activation leads to reduced plasticity of SST INs that is mediated by receptor internalisation

We have previously shown that aTBS induced LTP in SST INs can be prevented by acutely activating GABA_B_Rs^11^. However, our data suggests that application of baclofen leads to reduced LTP machinery from SST INs. To determine whether sustained activation of GABA_B_Rs shape the long-term plasticity landscape of SST INs, we first performed whole-cell patch-clamp recordings in *str. O/A* from adult male and female transgenic mice expressing YFP (Venus variant) under the vGAT promoter (Figure 5A). We recorded 108 *str. O/A* INs for the current study, those that were recovered (n=70) typically had horizontal dendrites confined to *str. Oriens* (*str. O*), with axons projecting to *Str. L-M* and were uniformly SST immunoreactive (Figure 5B). Recorded SST INs displayed prominent I_h_-mediated sag potentials, and possessed a medium to high rate of action potential (AP) discharge (Figure 5C), which was lower after baclofen pre-application; due to reduced input resistance and elevated rheobase (Supplementary Table 1).

**Figure 5:**
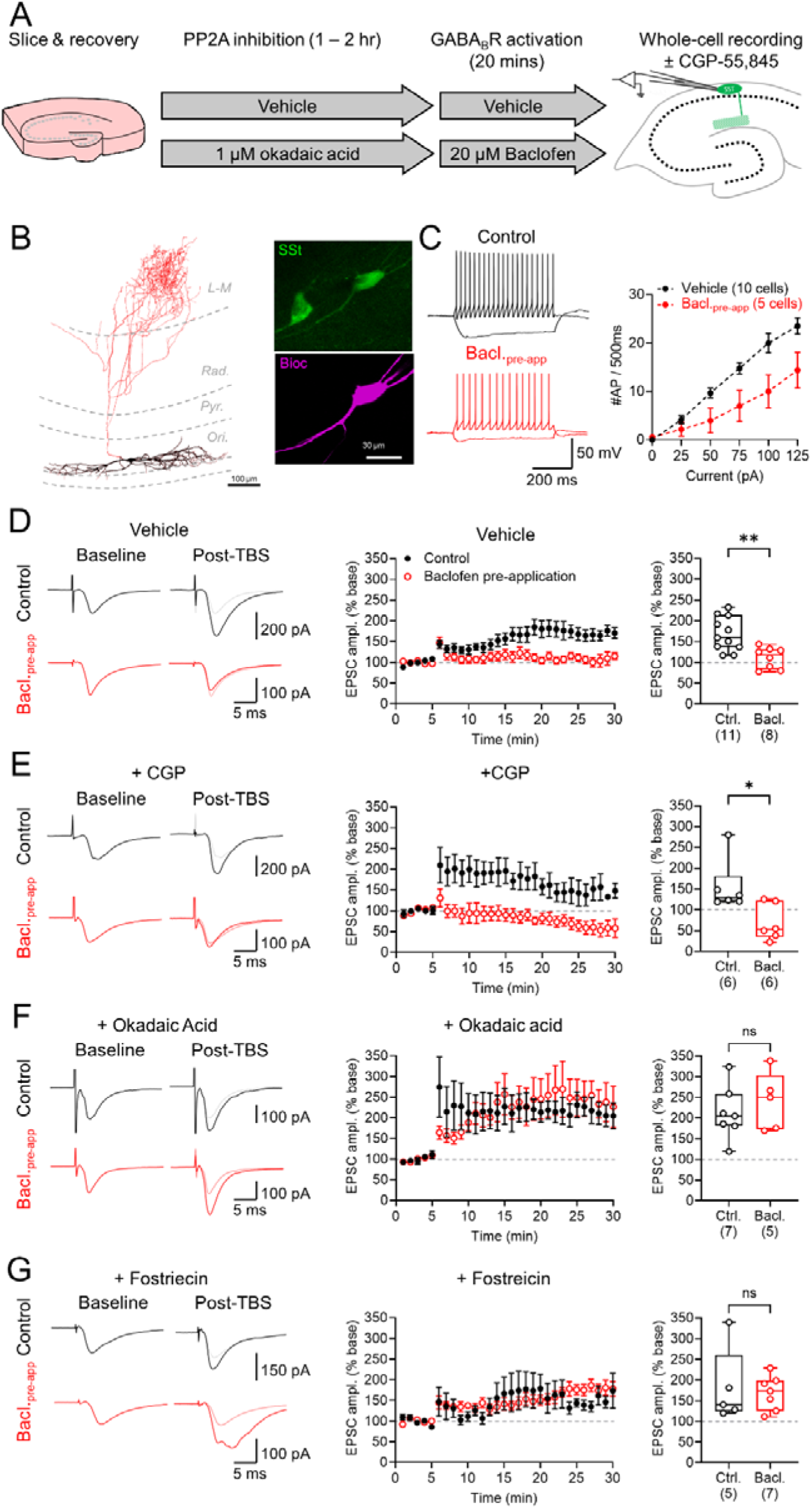
LTP of local inputs to SST INs is abolished by prolonged baclofen application. **A** Schematic of recording conditions, highlighting treatment periods and recording times. **B** Example reconstruction of a SST IN recorded from *str. O/A* of the adult mouse CA1 region, showing somatodendritic (black) and axonal (red) ramifications. Inset immunoreactivity for SST (green) and biocytin (purple). **C** Voltage responses in response hyper- to depolarising current steps (−500 to +500 pA, 100 pA steps, 500 ms duration) from SST INs recorded in following vehicle (black) and baclofen-pre-application (Bacl._pre-app_; red). Right, quantification of the action potential output of SST INs recorded under both conditions. **D** Example EPSC traces from vehicle (upper, black) and 20 μM baclofen pre-application (lower, red) measured under control conditions; for aTBS-LTP traces baseline EPSCs are shown underlain (grey and pink). Middle, time-course of plot of EPSC amplitude before and after aTBS-LTP induction for vehicle (black) and baclofen pre-application (red); 100% baseline level is shown for reference (grey dashed line). Right, bar-chart of EPSC amplitude change post aTBS-LTP (25-30 minutes) compared to baseline for vehicle (black) and baclofen pre-application (red); 100% baseline level indicated (grey dashed line). **E** aTBS-LTP recordings from SST INs performed in the presence of 5 μM CGP-55,845 (CGP) according to the same scheme as **D**. **F** aTBS-LTP following pre-application of slices with 1 μM okadaic acid for 1 hour with subsequent vehicle or baclofen pre-application. **G** aTBS-LTP in slices pre-application with 50 nM fostriecin for 1 hour prior to vehicle or baclofen pre-application. All data are shown as either mean ± SEM (time-course plots) or box-plots showing 25-75% box with median, with maximum range shown and individual cell results. Statistics shown as: ns – p>0.05, * - p<0.05, ** - p<0.01from Mann-Whitney non-parametric tests.

Under vehicle control conditions, we observed excitatory postsynaptic currents (EPSCs) resulting from *alveus* stimulation of 197.8 ± 17.6 pA (13.3 ± 3.9 V stimulation, n=11 SST INs), which following aTBS-LTP induction led to a facilitation of 68.5 ± 12.9% above baseline levels (p=0.001, Wilcoxon matched-pairs test). In baclofen pre-application slices, we observed baseline EPSCs with an average amplitude of 195.6 ± 54.6 pA (8.5 ± 2.9 V stimulation, n= 8 SST INs), which did not differ from vehicle recordings in amplitude (U_(11, 8)_= 28, P=0.21, Mann-Whitney) or stimulation strength (U_(11, 8)_= 33, P=0.38, Mann-Whitney). An aTBS-LTP induction in baclofen pre-application slices failed to increase EPSC amplitude above baseline levels (11.3 ± 9.6% above baseline, p=0.38, Wilcoxon matched-pairs test), and which was lower than for vehicle conditions (U_(11,8)_= 10; p=0.004, Mann-Whitney). To confirm that this loss of aTBS-LTP was not due to residual direct activation of GABA_B_Rs we performed the same experiments, but in which the GABA_B_R antagonist CGP-55,845 (CGP, 5 μM) was applied to the bath through-out recording (Figure 5E). In the presence of CGP, we still failed to induce aTBS-LTP in baclofen pre-application slices (31.3 ± 19.6% below baseline, p=0.31, Wilcoxon matched-pairs test), despite persistent induction in vehicle treated slices (54.5 ± 28.0% above baseline p=0.031, Wilcoxon matched-pairs test; U_(6,6)_=5; p=0.041 Mann Whitey); confirming that aTBS-LTP inhibition was not due to direct receptor activation.

As postsynaptic GABA_B_Rs undergo PP2A dependent internalisation (Figure 3) ^26,30,31,38^, we next asked if the impaired aTBS-LTP in SST INs was due to internalisation of the receptor. For these experiments, we incubated slices with OA (1 μM) for 60-120 minutes prior to vehicle or baclofen pre-application. Following OA incubation, we continued to observe reliable aTBS-LTP in both vehicle slices (112.1 ± 26.4% above baseline, p=0.016 Wilcoxon paired test) and baclofen pre-application slices in SST INs (140.1 ± 31.3% above baseline, p=0.029 Wilcoxon paired test), which were not different (U_(7,5)_ = 15; p=0.76, Mann-Whitney test; Figure 5F). As our SDS-FRL labelling was not possible with OA (Supplemetary Figure 3), we next confirmed that fostriecin (50 nM) produced similar effects to OA. After fostriecin incubation (60-90 minutes), we observed a tendency towards aTBS-LTP in vehicle slices (81.8 ± 45.6% above baseline, p=0.063 Wilcoxon paired test) and robust LTP in baclofen pre-application slices (69.0 ± 17.0% of baseline, p=0.016 Wilcoxon paired test), which was not different from vehicle (U_(5,7)_=16, p=0.876, Mann-Whitney test; Figure 5G).

Combined, these data indicate that sustained GABA_B_R activation impairs aTBS-LTP induction in SST INs, which is independent of direct receptor activation and which relies on internalisation of the receptor via PP2A-dependent phosphorylation.

### Prolonged activation of GABA_B_Rs impairs presynaptic release of GABA, but not postsynaptic expression of GABA_B_Rs in CA1 PCs

Assessing the wider function of GABA_B_R signalling following sustained pharmacological activation is critical to infer circuit wide effects. In CA1 PCs, following baclofen pre-application we observed a tendency towards reduced GABA_B1_, but unchanged Ca_V_1.2 surface expression. We next determined if these effects altered inhibitory signalling in and onto CA1 PCs. Therefore, we recorded GABA_B_R-mediated slow IPSCs in the presence of CNQX (10 μM), DL-AP5 (50 μM) and gabazine (10 μM) to block ionotropic receptor signalling (Figure 6A). We found that slow IPSCs were substantially reduced following baclofen pre-application (U_(10,7)_=3, p=0.0007, Mann-Whitney test; Figure 6B). However, baclofen-mediated whole-cell currents (I_WC_) were unaffected by baclofen pre-application (Figure 6C), with normal peak current-density (U_(9,7)_= 15.5, p=0.10, Mann-Whitney test; Figure 6D). One explanation for this disconnect between synaptic and pharmacological activation of GABA_B_Rs could be impaired GABA release. As such, we measured the input/output (I/O) relationship of ionotropic GABA_A_R-mediated IPSCs following *str. L-M* stimulation as a proxy for synaptic strength. We found that for the same stimulus, fast IPSCs were consistently lower in amplitude (Figure 6E), which upon quantification revealed 46% lower I/O slopes (U_(7,6)_=4, p=0.014, Mann-Whitney test, Figure 6F). To determine whether baclofen pre-application alters presynaptic GABA_B_R function, we next bath applied 10 μM baclofen to recordings of fast IPSCs (Figure 6G). Slices with pre-application of baclofen tended to have lower GABA_B_R presynaptic inhibition of IPSCs in *str. L-M* (U_(7,6)_=7, p=0.051, Mann-Whitney test, Figure 6H). Finally, we compared the effect of baclofen pre-application on CA1 PC intrinsic excitability. Contrary to what we observed in SST INs, we find that CA1 PCs are largely unaffected by baclofen pre-application, with subtle effects on membrane capacitance and action potential kinetics observed (Table S2).

**Figure 6:**
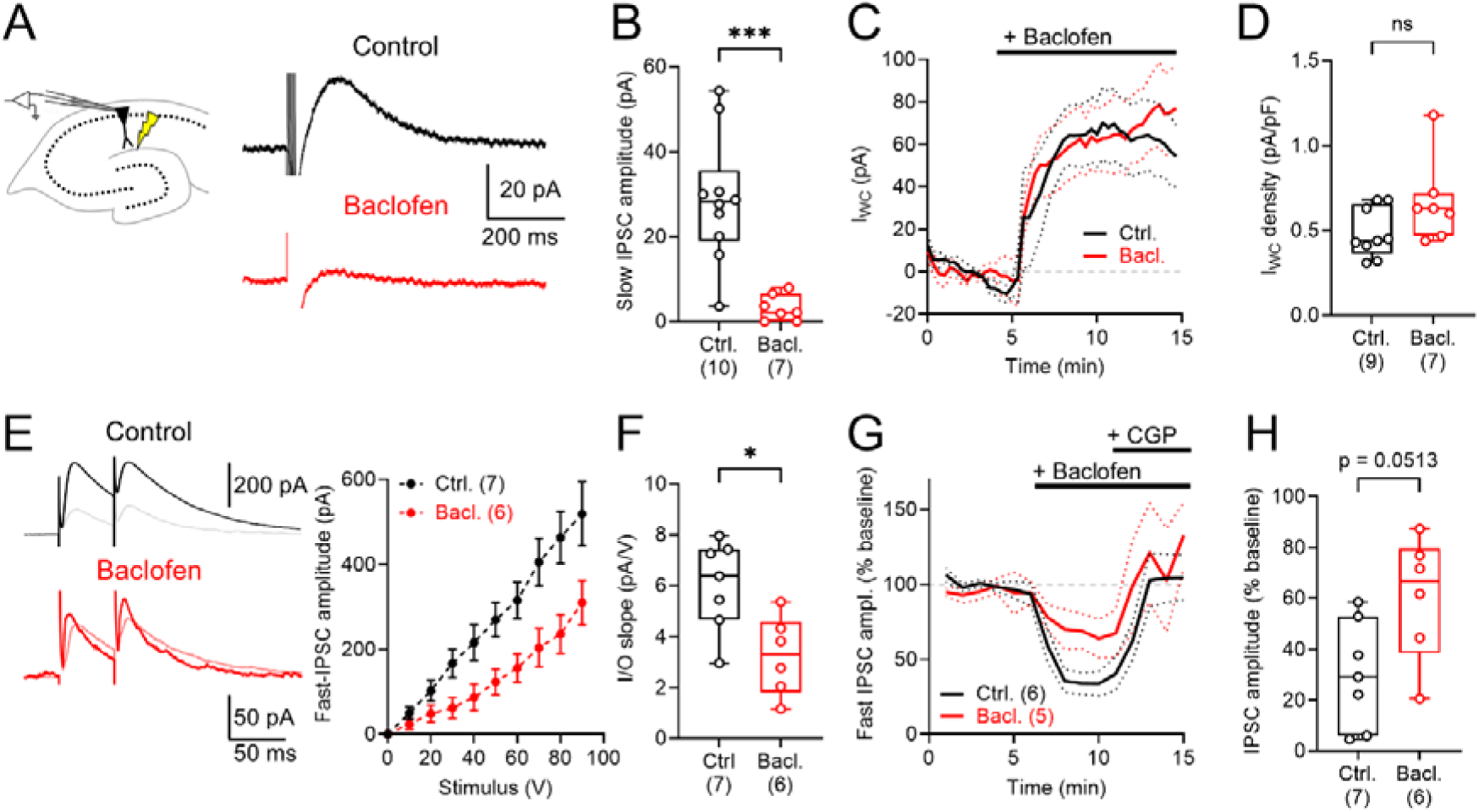
Prolonged GABA_B_R activation decreases GABA release in str. L-M, specific for presynaptic but not postsynaptic GABA_B_R-mediated inhibition. **A** Experimental schematic of recordings from CA1 PCs and example slow IPSCs recorded following 5x 200 Hz stimulation of *str. L-M* in the presence of CNQX (10 μM), DL-AP5 (50 μM) and gabazine (10 μM) from −65 mV voltage-clamp from control slices (black) and those pre-application with 20 μM baclofen (red). **B** quantification of slow IPSC amplitudes from control and baclofen pre-application slices. **C** Time course of whole-cell current (I_WC_) recorded from - 65 mV following bath application of 10 μM baclofen in control and baclofen pre-application slices. **D** Quantification of I_WC_ current density, normalised to cell capacitance. **E** Fast IPSCs evoked following paired stimulation of *str. L-M* from 0 mV using Cs-gluconate internal solution in the presence of CNQX (10 μM), DL-AP5 (50 μM) from control and baclofen pre-application slices. Traces recorded following bath application of 10 μM baclofen are shown underlain (grey/pink). Right, input-output (I/O) plot of fast IPSC amplitude from both groups. **F** Quantification of I/O slope for fast IPSCs. **G** Time course of fast IPSC amplitude before, and following bath application of 10 μM baclofen and 5 μM CGP-55,845 (CGP) in control and baclofen pre-application slices. **H** Quantification of baclofen mediated inhibition of fast IPSC amplitudes, expressed as % of baseline. Data is shown as either mean ± SEM (**C, E, G**) or box-plots showing 25-75% box with median, with maximum range (**B, D, F, H**). Individual cell data is shown overlaid. Statistics shown as: ns – p>0.05, * - p<0.05, *** - p<0.001; from Mann Whitney non-parametric tests.

Together these data suggest that prolonged GABA_B_R activation has limited effects on the postsynaptic function of CA1 PCs, in particular we find no evidence for reduced functional GABA_B_R expression. However, due to strong reductions in presynaptic GABA release, postsynaptic IPSCs are functionally reduced.

### GABA_B_R-dependent internalisation of LTP proteins in SST INs leads to enhanced temporoammonic plasticity onto CA1 PCs

SST INs are known to control the ability of CA1 PCs to undergo plasticity, notably in their distal dendrites in *str. L-M* aligned with entorhinal cortex inputs ^23^. To determine whether prolonged activation of GABA_B_Rs, and subsequent loss of SST IN plasticity alter temporoammonic LTP we performed extracellular field recordings from *str. L-M* of CA1, in which we induced plasticity at SST INs via repetitive stimulation of the alveus, followed by high frequency stimulation (HFS, 1× 100 Hz) to the temporoammonic pathway in vehicle or those with pre-application of 20 μM baclofen for 20 minutes (Figure 7A). Following HFS to *str. L-M* we observed increased fEPSP amplitudes when measured in all recordings from both treatment groups (Figure 7B), which when measured at 50-60 minutes post induction were 25.6 ± 7.8% above baseline in vehicle control recordings, and which did not differ in baclofen pre-application slices (20.5 ± 10.4%; U_(15,19)_=104, p=0.19, Mann-Whitney test, Figure 7C). However, we noted that a higher proportion of slices pre-application with baclofen failed to undergo LTP (>10% above baseline) as compared to vehicle controls (χ^2^=7.5, p=0.0062 Chi square test; Figure 7D). Therefore, we next analysed only those recordings in which LTP was observed (Figure 7E). Further quantification of successful LTP recordings (Figure 7F) revealed that post-tetanic potentiation (0-1 minute post HFS) was 34% higher in baclofen pre-application slices (U_(11,10)_= 25, p=0.018, Mann-Whitney test) and fEPSPs measured at 20-30 minutes (consistent with whole-cell recordings) were also 24% higher than control slices (U_(12,10)_=27, p=0.015, Mann-Whitney test), while LTP measured at 50-60 minutes was higher, albeit not significantly so (U_(12,10)_=37, p=0.07, Mann-Whitney test). We observed no difference in fEPSP slope relative to stimulus strength between control and baclofen pre-application slices (Figure 7G). These effects were likely specific to SST INs, as performing the same recordings but in the presence of the mGluR1α specific antagonist LY367,385 (100 μM), prevented any baclofen pre-application effects on temporoammonic LTP (Figure S4). This also excludes the effects of baclofen pre-application on GABAergic release and CA1 PC excitability as being causal to this effect.

**Figure 7:**
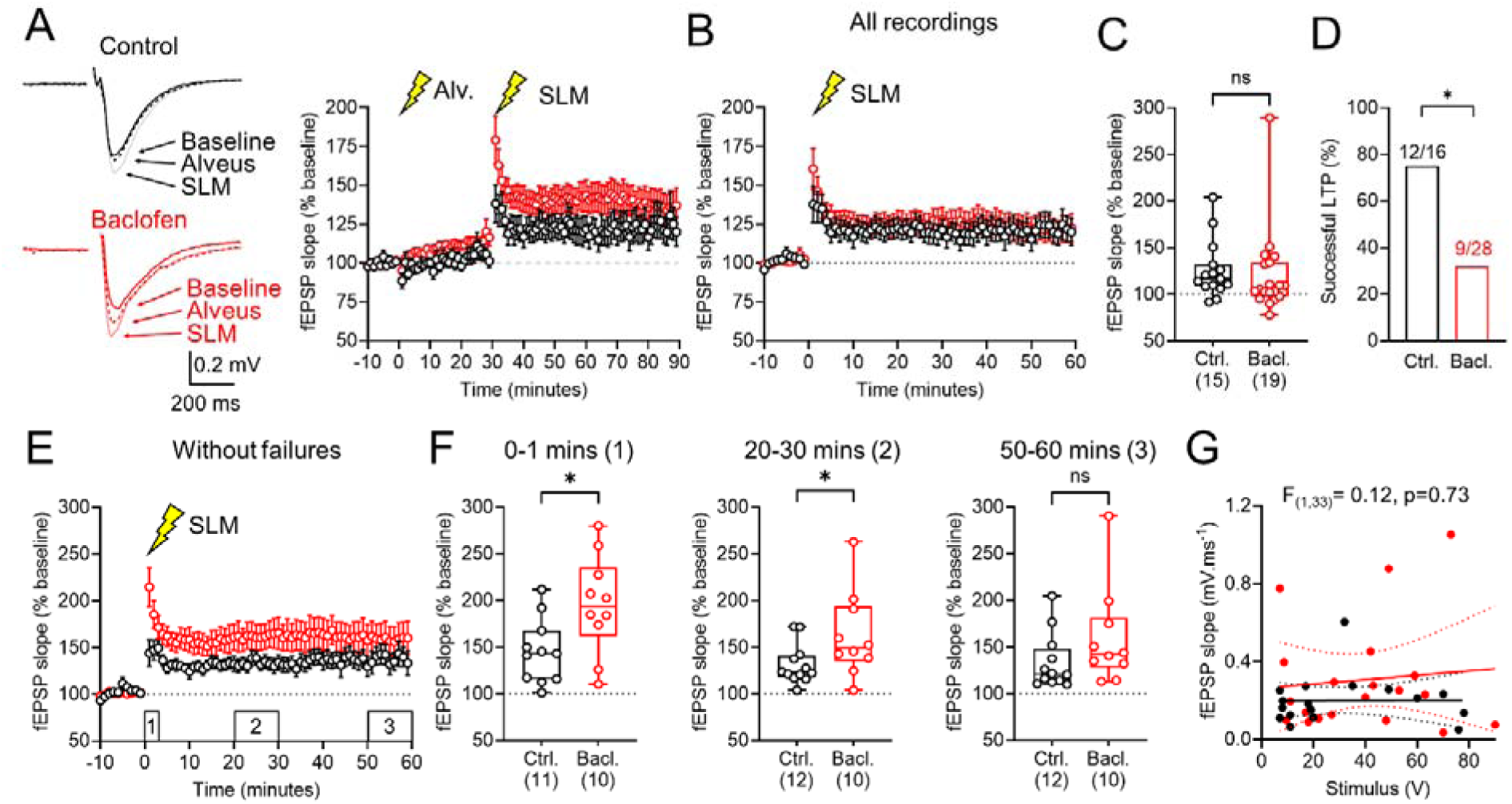
Prior GABA_B_R activation abolishes SST IN inhibition of short-term temporoammonic LTP. **A** Example fEPSP responses recorded in *str. L-M* following temporoammonic stimulation, from control (black) and 20 μM baclofen pre-application (red) slices. Traces are shown for initial baseline (solid line), following *alveus* LTP induction (dashed lines) and following *str. L-M* 1× 100 Hz HFS (grey/pink lines). Right, summary time-course of all fEPSP recordings normalised to the initial baseline, indicating LTP induction points (lightning bolts). **B** Time-course of LTP induced in *str. L-M* (SLM, lightning bolt), normalised to the 20-30 minutes baseline following *alveus* stimulation in all recordings. **C** Magnitude of fEPSP facilitation at 50-60 minutes post str. L-M HFS in all recordings. **D** Proportion of recordings in both control and baclofen pre-application that successfully induced LTP (>10% facilitation at 50-60 minutes post HFS). **E** Time-course of LTP induced in *str. L-M* normalised to the 20-30 minutes baseline following *alveus* stimulation in recordings where LTP was induced. **F** Quantification of fEPSP facilitation in control and baclofen pre-application slices at 0-1 (left), 20-30 (middle), and 50-60 (right) minutes post *str. L-M* HFS. **G** comparison of fEPSP slope and stimulus strength for all recordings in both groups. Data is shown as either mean ± SEM (**A, B, E**) or box-plots showing 25-75% box with median, with maximum range (**C, F**), proportions (**D**), or as individual data (**g**). Individual cell data is shown overlaid. Statistics shown as: ns – p>0.05, * - p<0.05; from Mann Whitney non-parametric tests.

These data confirm that prolonged GABA_B_R activation leading to a down-regulation in surface expression of GABA_B1_, mGluR1α, and Ca_V_1.2 in SST INs has profound effects on the ability of the CA1 microcircuit to encode inputs in *str. L-M*.

### Activation of GABA_B_Rs prevents acquisition of contextual fear memories

Finally, as SST IN plasticity has been causally linked to contextual fear conditioning when administered during training^39^, we asked if earlier baclofen administration impairs fear learning, For this, we administered mice with 2 mg/kg baclofen (or saline controls, both intraperitoneal injection) 1 hour prior to fear conditioning. Mice were then administered 3 foot-shocks (0.4 nA, 1-minute intervals, 5 minutes total) in the conditioning arena, and the proportion of time spent freezing measured. To determine contextual memory was retained by mice, they were returned to the same arena 24 hours later for 5 minutes (Figure 8A). Overall, saline control mice froze 42 ± 2% of the time during the initial test, increasing to 65 ± 4% freezing upon recall (p=0.004, Holm-Sidak test). By comparison, mice treated with baclofen 1 hour before training froze 17 ± 3% of the time, which did not increase upon recall (20 ± 7%, p=0.63, Holm-Sidak test), but which was markedly lower than saline controls (p<0.0001, Holm Sidak test; Figure 8B). These data confirm that prior administration of baclofen produces sustained impairments in contextual fear conditioning.

**Figure 8:**
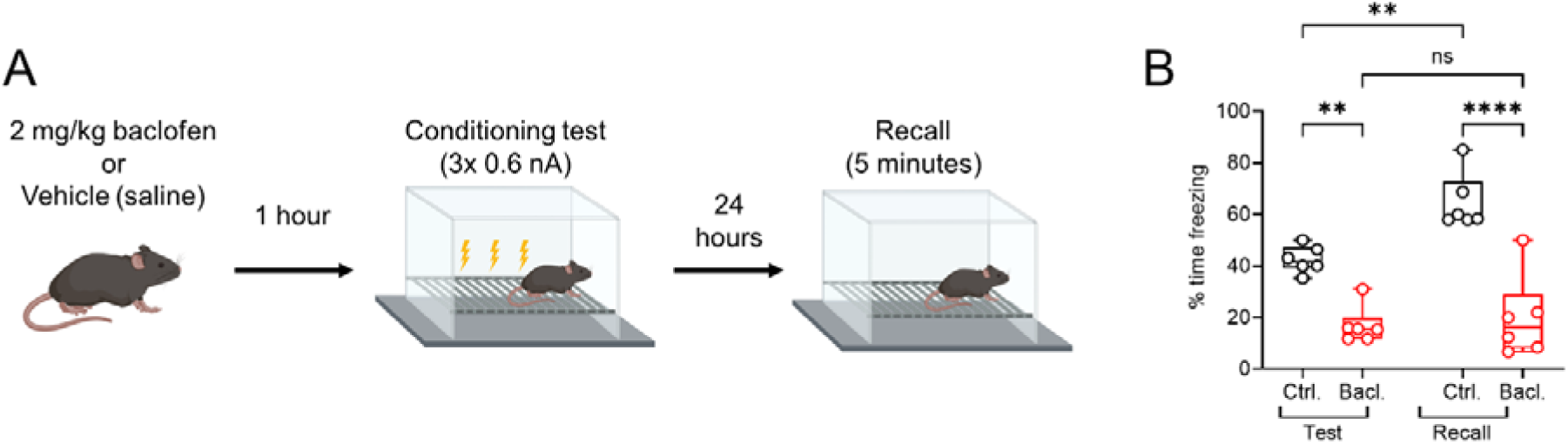
Prior GABA_B_R activation impairs contextual memory formation. **A** Schematic of contextual fear conditioning experiments, in which mice were dosed with 2 mg/kg baclofen 1 hour prior to training. **B** Quantification of % time freezing during the test and recall phases of conditional fear conditioning. Data is shown as box-plots showing 25-75% box with median, with maximum range. Individual cell data is shown overlaid. Statistics shown as: ns – p>0.05, ** - p<0.01, **** - p<0.0001; from 2-way ANOVA with Holm-Sidak post-tests.

This data suggests that prolonged GABA_B_R activation – as experience with in vivo administration - likely has lasting effects on the ability of hippocampal circuits to perform complex behavioural tasks and may provide important considerations for the use of baclofen therapeutically.

## Discussion

In this study we tested the hypothesis that sustained GABA_B_R activation leads to receptor internalisation in hippocampal SST INs, releasing inhibition of synaptic plasticity. We disproved this hypothesis and show baclofen administration leads to internalisation of GABA_B_Rs, Ca_V_1.2 and also mGluR1α in SST INs, which is mediated by PP2A-dependent phosphorylation. This internalisation impairs the long-term ability of SST INs to undergo synaptic plasticity, which in turn shifts the balance of plasticity at temporoammonic inputs and impairs contextual fear memory. These data reveal a novel mechanism of long-term disinhibition that has the potential to shape the plasticity landscape of the CA1 region over behaviourally relevant time-scales.

### PP2A-dependent GABA_B_R internalisation regulates SST IN activity

SST INs are a critical element in cortical circuits, gating the strength of inputs, and their plasticity, arriving from entorhinal cortex ^19^, and associative inputs within the hippocampus itself ^22,24^. These effects are mediated by a complex interplay of direct inhibitory actions on postsynaptic CA1 PCs, via presynaptic inhibition of glutamatergic and GABAergic inputs and through regulation of astrocyte function ^40^. Indeed, SST IN activity leads to profound disinhibition of the hippocampal circuit to strengthen CA3 inputs ^22^ and limit CA1 PC dendritic function ^24^. Thus, plasticity of SST INs themselves may act to shift the balance of CA1 function ^23^, which has a direct effect on CA3-dependent behaviours involving contextual information ^41^. Our work has shown that GABA_B_Rs acutely inhibit L-type VGCCs comprising Ca_V_1.2 subunits on SST INs to inhibit LTP ^11^, which led to our hypothesis that GABA_B_R internalisation would promote LTP induction, as this key inhibitory mechanism was lost. We have failed to prove this hypothesis, with the opposite appearing to be the case, due to coincident internalisation of the key LTP machinery, namely mGluR1α and L-type VGCCs; and which depends on activation of PP2A and that has been implicated in GABA_B_R phosphorylation in other brain areas ^26,30,31,38^.

Following GABA_B_R activation, adenylyl cyclase activity is inhibited, leading to lower cAMP levels and reduced PKA activation, which in turn limits the phosphorylation of Serine 892 on the GABA_B2_ subunit, which destabilises the receptor at the cell surface ^42^. Additionally, PP2A dephosphorylates Serine 783 on the GABA_B1_ subunit, which also destabilises the receptor leading to internalisation ^26^. mGluR1α is also a Type C G-protein coupled receptor (GPCR), thus it is highly plausible that these receptors are targeted for internalisation through the same intracellular cascade ^43^, particularly given PP2A has been shown to regulate recycling of mGluR1 isoforms ^44^. For Ca_V_1.2, the situation is more complex, as it is internalised upon GABA_B_R activation which appears to prevented by PP2A inhibition, while functional currents are not restored. Given the close proximity of GABA_B1_ to Ca_V_1.2 in SST IN dendrites^11^ it may form part of the GABA_B_R interactome in SST INs, as is the case for other Ca_V_ proteins^29^, thus function is impaired when GABA_B_Rs are lost. However, Ca_V_1.2 conductance is regulated by PKA phosphorylation^45^, thus, loss of GABA_B_R-mediated inhibition may counterbalance a numerical loss of channels. Beyond VGCCs, native GABA_B_Rs interact with a variety of transmembrane and cytosolic proteins in pyramidal cells ^28,29^, if the same interactome is present in SST INs and whether these proteins are similarly lost from these cells remain unexplored. Indeed, the functional association of GABA_B_Rs with other postsynaptic effectors (e.g. Kir3 channels) may differ in diverse INs ^8,9,12^. As INs make up only approximately 10% of neurons in the hippocampus ^46^, efforts to identify the GABA_B_R interactome using bulk protein samples^29^ have likely underestimated cell-type specific effects. Future recent advances in single-cell proteomic analysis ^47^ may allow detection of cell-type specific GABA_B_R interactomes, crucial to determine such function. Our data provides limited evidence of impaired postsynaptic GABA_B_R function in CA1 pyramidal cells under the same conditions, further suggesting cell-type specificity.

### GABA_B_R internalisation on SST INs impairs hippocampal circuit function

It has long been appreciated that acute GABA_B_R activation leads to paradoxical functions at the circuit level, notably enhanced LTP ^48,49^ and increase seizure activity^50^, despite directly hyperpolarising membranes ^51,52^ and inhibiting pre- and postsynaptic Ca^2+^ channels ^53,54^ on PCs. These effects point to a function beyond direct inhibition (favouring a disinhibitory model), exemplified by GABA_B_R inhibition on multiple IN types across the hippocampus ^4^. Our data extends this, providing evidence that prolonged GABA_B_R activation fundamentally shifts the ability of SST INs to undergo LTP on behaviourally relevant timescales. Given that SST IN LTP has been well associated with numerous behavioural functions, including spatial and contextual memory ^24,41,55–57^, this indicates a potential role of GABA_B_Rs in regulating this activity. We confirm that LTP in SST INs is crucial to maintain transfer of information to CA1 via the TA path, as previously reported ^19^, the control of which is impaired when baclofen is pre-applied. Furthermore, we show that GABA_B_R activation prior to contextual fear conditioning is sufficient to prevent memory encoding. While baclofen has been shown to impair fear conditioning previously ^39^, these studies relied on administering the drug immediately (<15 minutes) before fear acquisition, suggesting direct inhibitory effects. The pharmacokinetics of baclofen suggest peak brain concentrations with 30 minutes of IP administration, which remain high for several hours^58^. Therefore, it is nigh on impossible to estimate when (and if) loss of SST IN LTP supersedes inhibition driven by GABA_B_R inhibition. Future studies investigating GABA_B_R surface localisation on SST INs over hours and days post baclofen injection may reveal the temporal resolution of this relationship, but are outside the scope of this project.

### Translational importance

Baclofen is a common and important medicine, primarily prescribed for muscle spasm, but also tested for alcohol dependency, neurodevelopmental disorders, among other conditions (reviewed in ^59^), up to 100 mg/day; but is associated with many common neurological side effects. Thus, understanding how baclofen affects cortical circuits may inform its best use, particularly in conditions where GABAergic inhibition has also been implicated.

One of the most common clinical features associated with baclofen administration is the paradoxical induction of seizure activity, both in patients ^60^ and in rodent models ^50^. Originally proposed as a potential anti-seizure medication ^61^, effects of baclofen on hippocampal circuits has previously been described, leading to seizure like activity ^62^. The baclofen-dependent loss of SST IN excitatory signalling we observe may favour long-term reduction in local SST IN inhibition, which would be predicted to cause seizures ^63^. Interestingly, we recently showed higher GABA_B_R signalling in PCs in patients who have experienced seizures ^64^. It is tempting to speculate that such increases may result from reduced inhibitory tone, leading to compensatory upregulation of receptor expression. Further study is required to determine if this is the case.

A recent off-label use for high-dose baclofen has been in treating addiction, particularly alcoholism use disorder ^65^. In rodents, GABA_B_Rs have been shown to be lost in the cortex following chronic alcohol administration ^66^ and ventral tegmental area (VTA) following administration of amphetamine or cocaine ^26^, albeit not tested in SST INs. However, it has been shown that acute alcohol administration bidirectionally affects SST INs, with low doses increasing activity and high doses attenuating activity ^67^. SST INs are present in cortical ^15,68^ and VTA ^69^ circuits, and their activity is strongly downregulated in chronic alcohol consuming mice, which may be corrected by inhibiting them directly^70^. Our data suggests that prolonged baclofen administration may prevent SST IN activity, akin to chemogenetic approaches used previously, thus may reflect a mechanism by which baclofen could produce therapeutic benefit.

### Limitations

We have identified some key limitations of our current study. First, while we show that internalisation of GABA_B_Rs from SST IN dendritic membranes occurs following 20 minutes of baclofen application, using our approach we were not able to identify the specific temporal dynamics of this loss. If such effects occur rapidly (within seconds) or require longer (minutes to hours) remains unclear. Nevertheless, baclofen as a therapy is often administered at moderate to high doses (up to 1g/day) and has a CSF half-life of several hours ^58^, thus such temporal dynamics – whilst mechanistically interesting – are somewhat negated by these pharmacokinetics. Further study identifying whether GABA_B_R internalisation on SST INs displays dose- and time-dependence would provide useful insight to baclofen’s therapeutic use. Furthermore, we have not identified how long it takes SST INs to recover LTP following baclofen administration, which may also provide important therapeutic information.

One unusual observation in our data was that okadaic acid appeared to occlude labelling of GABA_B1_ in SDS-FRL experiments, both on SST INs and CA1 PCs (Figure S3). We do not have a good explanation for this effect. One possibility is that SDS-FRL relies on immunolabelling of receptors ^71^ and okadaic acid as a large polypeptide interacts with an epitope on PP2A close to its interaction site with GABA_B1_. Such an interaction may mask or occlude antibody binding, that would otherwise not be observed with the use of small molecule pharmacology, such as fostriecin. A similar phenomenon has been proposed for the binding of the amyloid precursor protein (APP) to GABA_B1_, which has been indicated in SDS-FRL to be diminished in APP overexpressing mice^72^, but in which mice GABA_B_R function is unaltered ^73^. These data indicate the importance of providing multiple lines of evidence to support conclusions based on physiology or anatomy alone, which we have provided through comparative use of okadaic acid and fostriecin that both selectively inhibit PP2A ^74^.

Finally, our data suggest that in SST INs GABA_B_Rs likely interact with themselves, and influence the function of – mGluR1α, Ca_v_1.2, and PP2A. Such proteins do not appear on previously published interactomes of GABA_B1_ or GABA_B2_ in the hippocampus ^29^. Our data suggest the existence of cell-type-specific interactomes, which may not be detectable at a whole brain, or even region-specific level. Such an endeavour would almost certainly identify cell-type specific differences even within neurochemically defined classes, as highlighted by previous work ^4,12^. However, such analyses are beyond the scope of the current study. Future work should seek to identify which proteins GABA_B_Rs interact with, in a cell-type specific manner, and potentially if such interactions change with age or neuropathology.

### Summary and outlook

Here we show in hippocampal SST INs that prolonged activation of GABA_B_Rs is sufficient to cause internalisation of the receptor through PP2A-dependent mechanisms, which also leads to loss of mGluR1α and Ca_V_1.2 to prevent synaptic plasticity at CA1 PC inputs. These data provide compelling evidence of how the inhibitory molecule baclofen exerts a paradoxical disinhibitory effect by reducing SST IN function, which has relevance for hippocampal function and disease.

## Acknowledgements

We wish to thank Natalie Wernet and Julia Bank for technical support, BVS technicians for supporting animal husbandy, and members of the Booker, Vida, and Kulik labs for discussions of the data. Funding was provided by: UK Medical Research Council (MR/Y014529/1, SAB); DFG (FOR 2134, IV, AK), BIOSS-2 (AK), CRC-TRR 384/1 (IV, AK) the Patrick Wild Centre (SAB), and the Simons Initiative for the Developing Brain (SFARI: 529085 - SAB)

## Author Contributions

Conceptualisation: SAB, AK; Methodology: NS, SAB, MW, AS, DL, AK; Validation: SAB, AK; Formal Analysis: SAB, AK, NS, MW, AS; Investigation: SAB, NS, MW, DL, AS, RL; Writing original draft: SAB; Writing – review and editing: SAB, NS, IV, AK; Supervision: SAB, IV, AK; Funding acquisition: SAB, IV, AK.

## Materials and methods

### Animals

All *ex vivo* electrophysiology was performed in brain slices prepared from male and female mice (2-4 months old), with all procedures performed according to Home Office (ASPA, 2013) and The University of Edinburgh Ethical Board guidelines. Mice were maintained on a C57/Bl6J^CRL^ background and expressed yellow-fluorescent protein (YFP) under the vesicular GABA transporter (vGAT)^75^. Mice were housed on a 12h light/dark cycle, with *ad libitum* access to food and water.

For SDS-FRL, male mice (8-week-old, n=12) either wild-type or in which SST-INs selectively expressed Channelrhodopsin2(ChR2)-YFP fusion protein were derived from crossing SST-Cre and Ai32^RCL-ChR2(H134R)/EYFP^ transgenic mice. These mice and male Wistar rats (8-week-old, n = 3) were used for quatitative immuno electron microscopic analysis. For behavioural analysis, male C57/Bl6J mice (8–12 weeks old, n = 6 per group) were used. Care and handling of the animals prior to and during the experimental procedures followed European Union and national regulations (German Animal Welfare Act) and all experiments were performed in accordance with institutional guidelines (University of Freiburg, Germany) with permission from local authority (Freiburg, X21/04B).

### Acute brain slice preparation

Brain slices were prepared as previously described ^76^. Mice were terminally anaesthetised with isoflurane, decapitated, then their brains rapidly dissected into ice-cold sucrose-modified artificial cerebrospinal fluid (sucrose-ACSF; ACSF; in mM: 87 NaCl, 2.5 KCl, 25 NaHCO_3_, 1.25 NaH_2_PO_4_, 25 glucose, 75 sucrose, 7 MgCl_2_, 0.5 CaCl_2_) which was saturated with carbogen (95% O_2_/5% CO_2_). Brains were glued to a vibratome stage (Leica VT1200S, Leica, Germany), then 300 µm (for whole-cell recordings) or 500 µm (for extracellular recordings) horizontal slices containing the hippocampi were prepared. Following slicing, brain slices were placed in either: a submerged holding chamber containing sucrose-ACSF warmed to 35 °C for 30 min, then at room temperature; or on small squares of filter paper placed in a liquid/gas interface chamber, containing recording ACSF (in mM: 125 NaCl, 2.5 KCl, 25 NaHCO_3_, 1.25 NaH_2_PO_4_, 25 glucose, 1 MgCl_2_, 2 CaCl_2_) and bubbled with carbogen.

### Whole-cell patch clamp recordings

Briefly, 400 µm horizontal hippocampal slices were prepared from 60-90 day-old C57/Bl6J mice as previously described^77^. For recording, slices were transferred to a submerged chamber maintained at 31 ± 1°C. Whole-cell recordings were made using pipettes filled with either K-gluconate or Cs-gluconate based solutions. Voltage-clamp recordings were performed from a potential of either −65 mV or −60 mV, while current-clamp recordings were made from resting membrane potential. Intrinsic properties of recorded neurons were characterized on their voltage response to hyper- to depolarizing current steps, in current-clamp. HVA calcium transients, mGluR EPSCs, and GABA_B_R-mediated currents were measured in the presence of 10 µM CNQX, 50 µM DL-AP-5, and 50 µM picrotoxin. GABA_A_R IPSCs were elicited by placing a bipolar stimulating electrode in str. lacunosum-moleculare approximately 200-300 µm rostral to the recorded neuron, in the presence of DL-10 µM CNQX, 50 µM DL-AP-5. LTP was induced at alveus inputs to SST INs with EPSCs elicited by a bipolar stimulating electrode placed in the alveus, in the presence of 50 µM picrotoxin. Following baseline, associative LTP was induced by theta-burst stimulation paired with a depolarization to −20 mV, repeated 3 times at 30 second intervals. EPSCs were measured for 25 minutes following induction. For all stimulation, the pulse duration was 200 µs delivered via a constant voltage generator. For HVA VGCC activation, once a whole-cell patch-clamp recording was obtained using a Cs-Gluconate internal solution (containing QX-314), the cell was allowed to dialyse for 5 minutes. Then at −60 mV voltage-clamp, 3 families of depolarizing steps were applied (−60 to 20 mV, 5 mV steps, 500 ms duration). The L-type VGCC blocker nifedipine (20 µM) was then applied to the bath for 3 minutes and 3 further families of depolarizing steps obtained. All traces were then P/N subtracted based on the 5 mV step and the peak current measured. The peak current under nifedipine was then subtracted from control. For mGluR EPSC experiments, slow-EPSCs were elicited by focal puff application of the Group 1 mGluR agonist DPHG (100 µM, 500 ms, 20 psi, diluted from stock in 150 mM NaCl) delivered via a patch pipette placed ∼50 µm from the cell body. The specificity of the mGluR current was assessed by subsequent bath application of the mGluR1α-specific antagonist LY-367,385 (100 µM).

For pre-application experiments, slices were transferred to a holding chamber containing recording ACSF and 20 µM R-baclofen for 20 minutes, slices were then placed in the recording chamber and rinsed with ACSF without R-baclofen for 4-6 minutes prior to recording. To block PP2A activity, slices were transferred to a chamber containing ACSF and 1 µM Okadaic Acid for at least 3 hours prior to recording or baclofen pre-application. All other drugs were applied directly to the perfusing ACSF.

### Field recordings

For field excitatory postsynaptic potential (fEPSP) recordings, hippocampal slices were transferred to an interface recording chamber perfused with carbogenated artificial cerebrospinal fluid (ACSF) at 2–3 mL/min and maintained at 30 ± 1 °C. Recording electrodes (1–3 MΩ) were pulled from borosilicate glass capillaries (1.5 mm outer / 0.86 mm inner diameter; Harvard Apparatus, UK) using a horizontal puller and filled with recording ACSF. Slices were visualised with a wide-field microscope (Leica, Germany), and electrodes were positioned in the str. L-M of CA1.

Extracellular fEPSPs were evoked using a paired-pulse protocol delivered via a bipolar stimulating electrode placed in *str. L-M*, approximately 500 µm to 1 mm from the recording electrode, to activate the temporoammonic pathway. A second stimulating electrode was positioned in parallel within the alveus of CA1. To activate GABA¬B receptors, slices were perfused with 20 µM baclofen for 20 min, followed by a 20-min washout period.

A 10-min baseline was recorded under control conditions or following baclofen washout. LTP was first induced via alveus stimulation using a high-frequency stimulation (HFS) protocol consisting of three 100 Hz trains of 20 pulses delivered 30 s apart, or a protocol consisting of five 100 Hz trains of 5 pulses, delivered 30 s apart. LTP was then induced in str. L-M using an HFS protocol of 1× 100 Hz train of 100 pulses delivered 30 s apart.

Potentiation was monitored for 60 min following induction. LTP magnitude was quantified as the mean fEPSP slope measured 50–60 min post-induction, normalised to the 10-min baseline. LTP was considered successful if the 50–60 min slope exceeded baseline by >10%. Recordings were excluded if baseline stability, assessed by comparing mean slopes at 1–2 min and 9–10 min, varied ±10%. Signals were filtered online (1 Hz high-pass, 500 Hz low-pass) and digitised at 10 kHz. Data were acquired and analysed offline using WinLTP (v3.01, University of Bristol, UK).

### Neuronal visualization and immunohistochemistry

All neurons were filled with biocytin during recording, fixed overnight in 4% PFA and labelled with streptavidin and antibodies to SST according to previous methods^78^. Slices were washed in PBS then blocked for 1 hour at room temperature (10% NGS, 0.3% Triton X-100 and 0.05% NaN_3_ in PBS). Following blocking slices were incubated with primary antibodies at 4°C for 48h (1:1000 rabbit SST-14, T-4103, BMA Biomedicals, Switzerland; 5% NGS, 0.3% Triton X-100 and 0.05% NaN_3_ in PBS), washed in PBS then transferred to a secondary antibody solution containing fluorescent-conjugated streptavidin (1:500 goat anti-rabbit AlexaFluor 488, 1:500 Streptavidin AlexaFluor 633, 3% NGS, 0.1% Triton X-100, 0.05% NaN_3_). Following labeling the slices were washed in 0.1M PB and mounted on glass slides with Vectashield HardSet mounting medium (Vector Labs, UK). To identify SST Ins Confocal image stacks of recorded neurons were taken on a Leica SP8 confocal microscope using a 63x (1.4 NA) objective at 1024 × 1024 resolution (Z step size 1 µm).

### Electron microscopy

#### Acute slice preparation and pharmacology for SDS-FRL

Acute hippocampal slices were prepared as previously described ^76^. Animals were sedated with isoflurane then they were decapitated and the brains were rapidly dissected and chilled in semi-frozen carbogenated (95% O2/5% CO2) sucrose-substituted artificial cerebrospinal fluid (sucrose-ACSF, in mM: 87 NaCl, 2.5 KCl, 25 NaHCO3, 1.25 NaH2PO4, 25 glucose, 75 sucrose, 7 MgCl2, 0.5 CaCl2, 1 Na-pyruvate, 1 Na-ascorbate). Transverse hippocampal slices (200 µm thick) were cut on a vibratome (VT1200s, Leica, Germany) and stored submerged in sucrose-ACSF warmed to 35°C for at least 30 min and subsequently at room temperature (RT). Both solutions were equilibrated with 95% O2 and 5% CO2 gas mixture throughout experiments. Acute slices were divided into four groups for pharmacological treatment: 1, incubated in ACSF (control, 30 min), 2, ACSF + GABA_B_R agonist baclofen (Bac, 20 µM, 20 min), 3, ACSF + protein phosphatase 2A inhibitor fostriecin (FS, 50 nM, 90 min), 4, ACSF + fostriecin + baclofen (FS/Bac, FS 50 nM 90 min + Bac 20 µM, 20 min). After pharmacological manipulations, slices were transferred into fixative containing 1% paraformaldehyde and 15% saturated picric acid made up in 0.1 M phosphate buffer (PB) overnight (O/N) at 4 °C.

##### SDS-FRL immuneoelctron microscopy

After fixation, slices were cryoprotected in 30% glycerol in 0.1M PB O/N at 4°C ^10^. Blocks containing stratum oriens-alveus were trimmed from the slices and frozen under high-pressure (HPM 100, Leica). The frozen samples were fractured at −140°C and the fractured faces were coated with carbon, (5 nm), platinum-carbon (2 nm) and carbon (18 nm) in a freeze-fracture replica machine (ACE 900, Leica). Replicas were digested at 80°C in a solution containing 2.5% SDS and 20% sucrose diluted in 15 mM Tris buffer (TB, pH 8.3) for 20 hr. Subsequently, replicas were washed in washing buffer containing 0.05% bovine serum albumin (BSA, Roth, Germany) and 0.1% Tween 20 in 50 mM Tris-buffered saline (TBS) and then blocked in a solution containing 5% BSA and 0.1% Tween 20 in TBS for 1 hr at RT. Afterwards, replicas prepared from SST-cre mice were incubated in the following mixtures of primary antibodies in a solution containing 1% BSA and 0.1% Tween 20 made up in TBS for 3 days at 15°C: (i) GABA_B1_ (B17, rabbit, 10 µg/ml; Kulik et al., 2002) and green fluorescence protein (GFP-1010, chicken 0.4 µg/ml, Aves Labs, Oregon; Booker et al., 2020) or (ii) Ca_v_1.2 (mouse, 50 μg/ml, NeuroMab Facility, California; Booker et al., 2018) and GFP. Replicas were washed then reacted with 6 nm (GABA_B1_ and Ca_v_1.2) and 18 nm (GFP) gold particle-conjugated secondary antibodies (1:30, Jackson ImmunoResearch Europe, Cambridgeshire) O/N at 15°C. The replicas prepared from WT mice were incubated in Ca_v_1.2 (mouse, 50 μg/ml, NeuroMab Facility, California), GABAB1 (B17, rabbit, 10 µg/ml) and metabotropic glutamate receptor 1α-subunit (mGluR1α, Guinea pig, 0.4 µg/ml, Frontier Institute, Hokkaido; Booker et al., 2018) in a solution containing 1% BSA and 0.1% Tween 20 made up in TBS for 3 days at 15°C. Replicas were washed then reacted with 5 nm (GABA_B1_) 10nm (Ca_v_1.2) and 18 nm (mGluR1α) gold particle-conjugated secondary antibodies (1:30, Jackson ImmunoResearch Europe, Cambridgeshire) O/N at 15°C. Finally, replicas were washed in TBS then distilled water and mounted on 100-mesh grids.

#### Electron microscopy and quantitative analysis of protein density

Labeled replicas were analyzed under an electron microscope (JEM 2100 Plus). Since all antibodies target intracellular protein epitopes, immunoreactivity can be detected on the plasma membrane’s protoplasmic face (P-face) but not the exoplasmic face (E-face). The surface density of receptors and ion channels was determined by dividing the absolute number of particles labeling molecules of interest by the surface of the respective segments of the dendritic shafts of GFP-positive SST-INs on replicas.

### Behavioural analysis

To investigate the role of GABA_B_R activation in fear learning, we performed a fear conditioning experiment. Male C57/Bl6J mice (8–12 weeks old, n = 6 per group) were handled and habituated. On the day of the test, mice were given a two-minute baseline period before receiving three-foot shocks (0.06mA) spaced one minute apart. Following the last shock, the mice remained in the fear conditioning arena for an additional minute, then they were transferred to the home cage. The entire training session lasted for approx. five minutes. Mice were divided into test and control groups. Test group were injected with baclofen (i.p., 2 mg/kg), while the control group received a vehicle (0.9% NaCl). One hour following the baclofen or vehicle injection, freezing time was measured in the same context as recall in order to access fear memory.

### Statistical analysis

For *ex vivo* electrophysiology, one or two cells or slices were recorded per animal per treatment, limiting the ability to assess intra-animal variability. As such cells and slices were treated as independent replicates and statistical analyses were conducted on these data. Parametric and non-parametric tests were applied as appropriate, including two-way ANOVA, Student’s t-test, Mann–Whitney U-tests, and Wilcoxon signed-rank tests. For SDS-FRL, where many dendritic profiles were sampled per animal replicate, we performed linear-mixed effects modelling (LMM) of using the “lmer” function to limit the potential inclusion of pseudoreplication. Animal was assigned as random effects, then type III ANOVA (Satterthwaites method) was performed on the model to assess main effects (e.g. drug treatment). If a statistical interaction between main effects was observed, then post-hoc analysis was performed (Tukey post hoc tests). For all comparisons data is shown as box-plots with median and 25–75% percentiles displayed in the box; minimum and maximum extents are shown as whiskers, with individual data points shown from cell or animal averages. For timecourses and current-voltage plots, data is shown as mean ± SEM. Statistical significance was defined as p < 0.05. Statistical tests and graphing were performed using either GraphPad Prism (GraphPad Software v10.4.1, San Diego, CA, United States) or R-Studio.

