## Supplementary data for "PP2A-dependent internalisation of GABA_B_ receptors in somatostatin interneurons regulates function and plasticity"

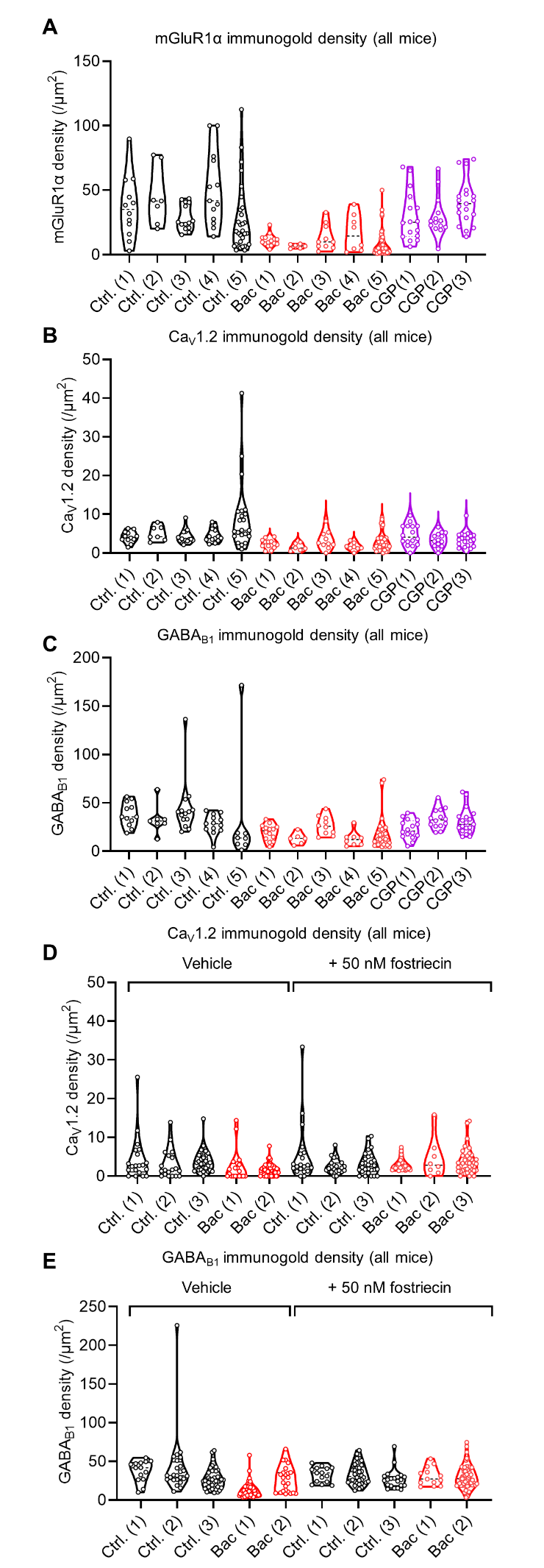

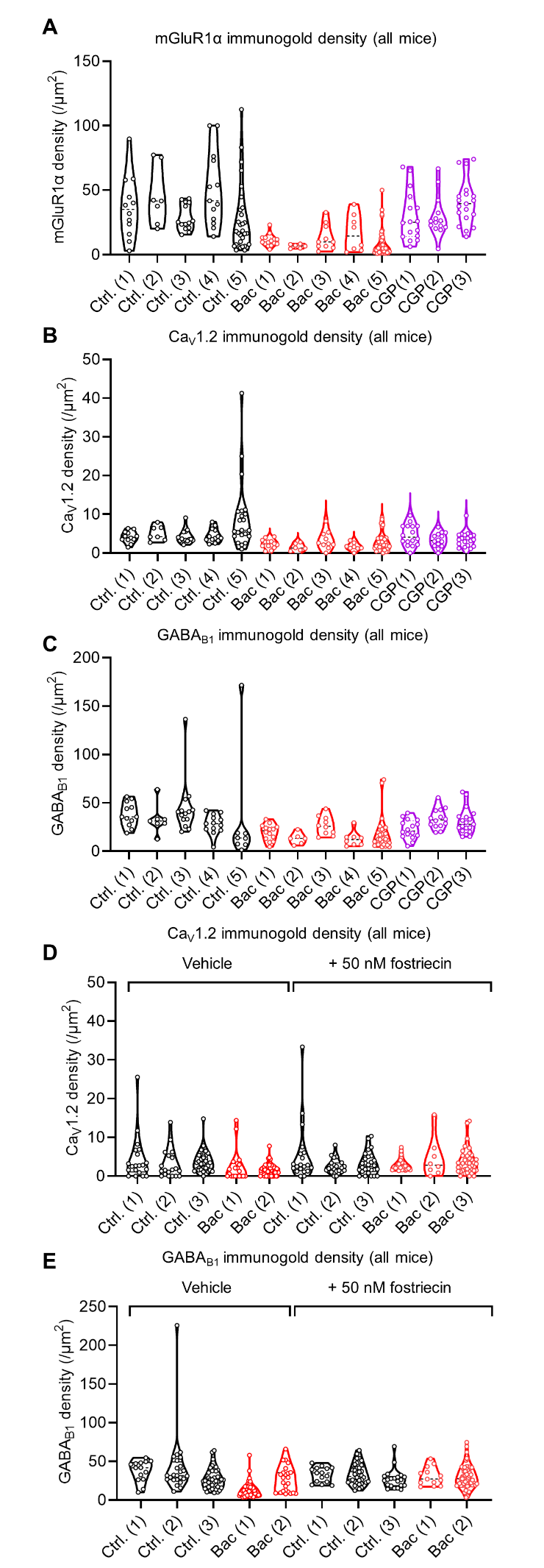

*Figure S1: Summary of all immunogold density measurements from all dendritic profiles measured in each biological replicate for the current study*. **A**) mGluR1α density in from SDS-FRIL replicas from slices treated with vehicle control (black), 20 minutes of 20 μM baclofen (red) or 5 μM CGP-55,845 (purple). **B**) the same data, but for Ca_V_1.2 labelling. **C**) the same data but for GABAB1 labelling. **D**) Ca_V_1.2 immunogold labelling on dendrites in SDS-FRIL replicas from mice expressing SST-ChR2 for control (black) and baclofen (red) treated slices. **E**) the same data as D but for GABAB immunogold labelling. All data is shown as individual density for each dendritic profile, with violin plots for each animal.

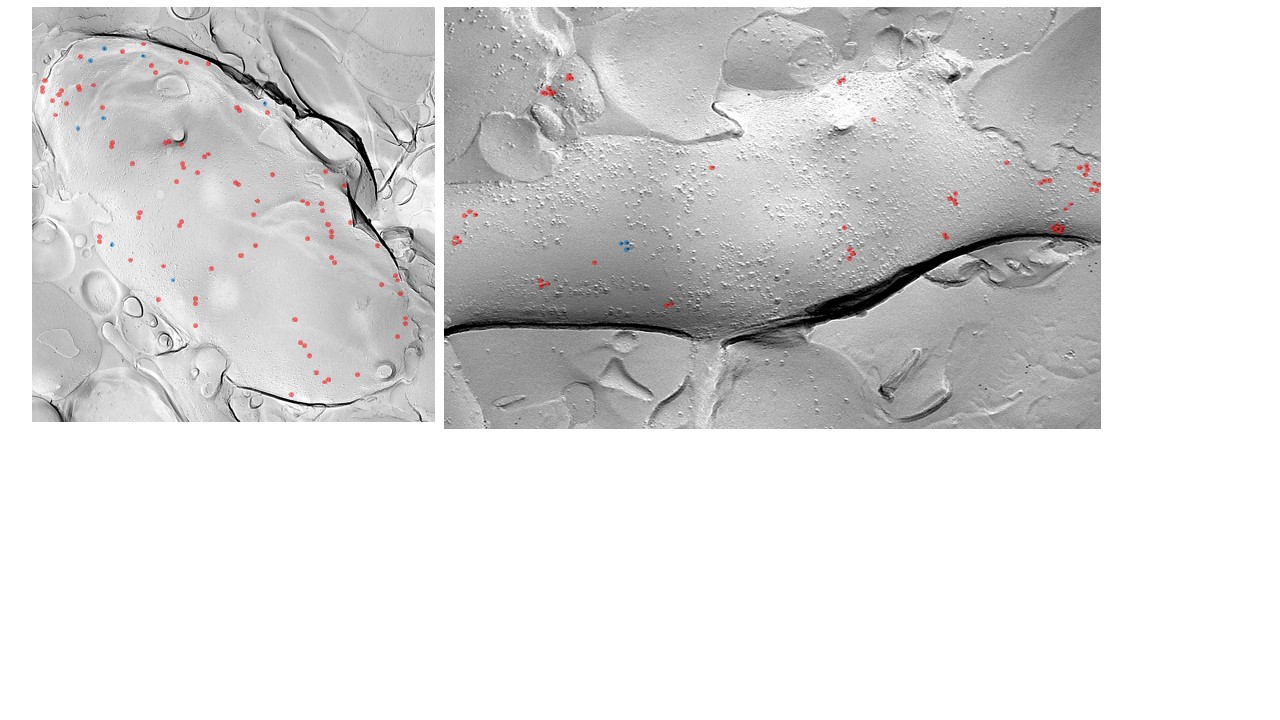

Cav1.2(10nm)

GABA_B1_ (5nm)

**Baclofen**

**Control**

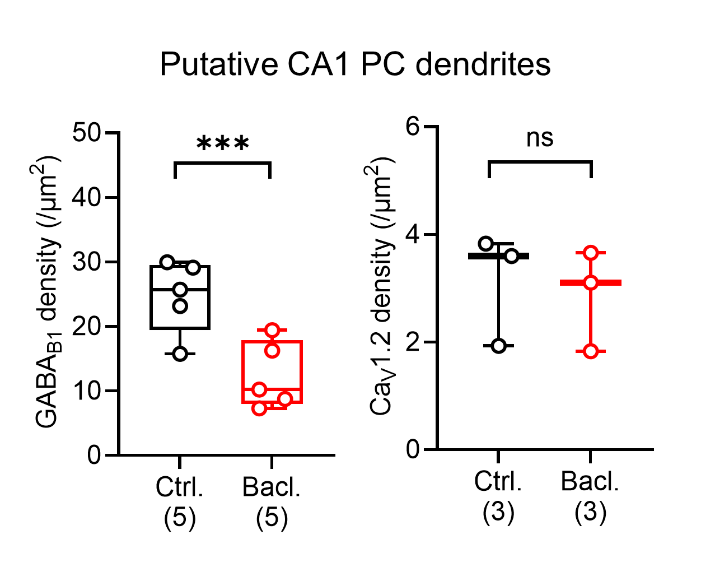

*Figure S2: Immunogold labelling for GABA_B1_ and Ca_V_1.2 on putative PC dendrites identified in the same replicas as those shown in Figure 1.* Putative CA1 PC dendrites were identified on the basis of possessing dendritic spines a prominent postsynaptic density. Data is shown from control (black) and baclofen pre-treated (red) replicas from 3 mice, and depicted as box-plots overlaid by individual animal averages. Statistics shown from LMM analysis, ns – p>0.05, *** - p<0.0001.

MGluR1α (18nm)

Cav 1.2 (6nm)

MGluR1α (18nm)

GABA_B1_ (6nm)

**Okadaic acid treated SST den**

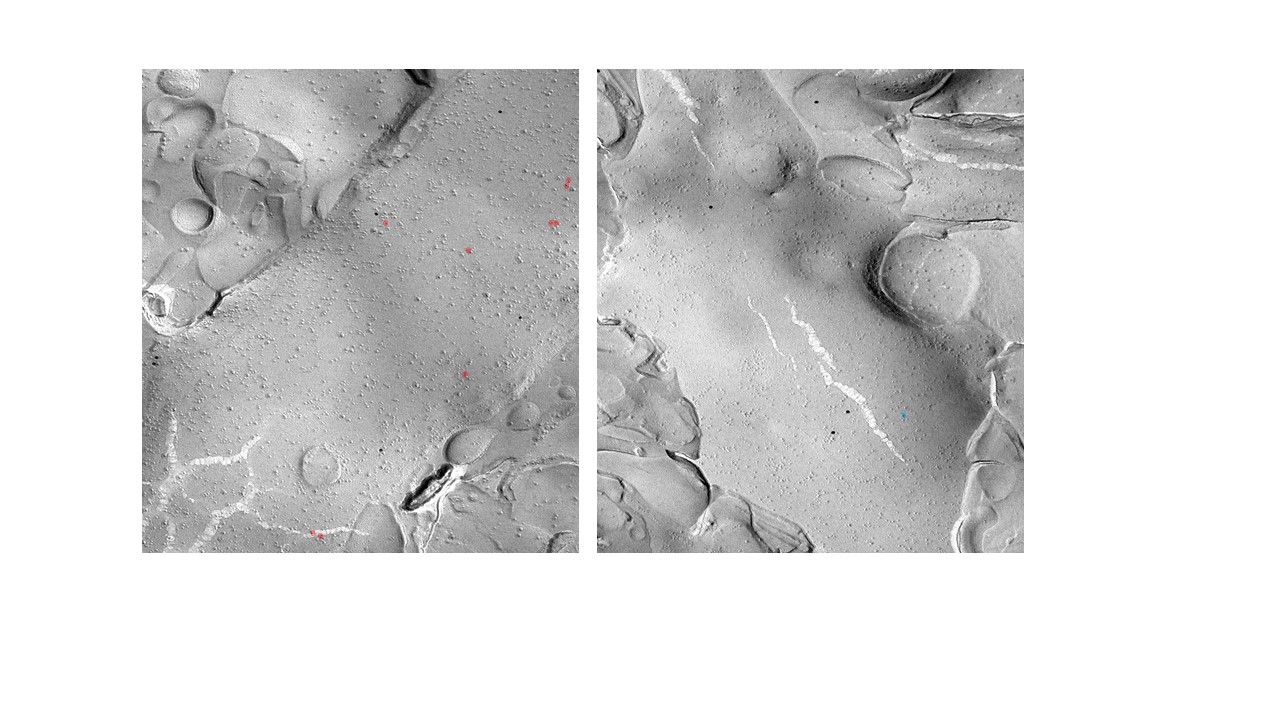

Cav 1.2 (6nm)

GABA_B1_ (6nm)

**Okadaic acid treated PC den**
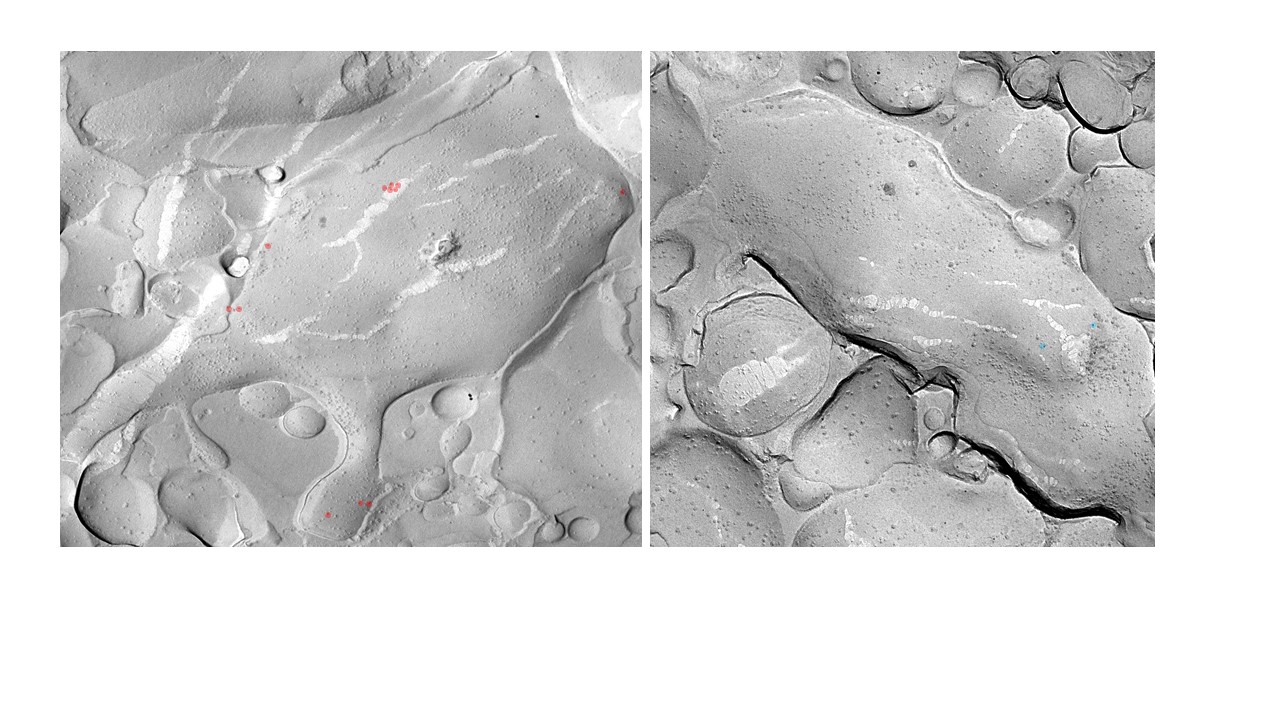

*Figure S3: Okadaic acid prevents occludes GABA_B1_ immunolabelling in SDS-FRL*. Example images from a putative SST IN (top) and CA1 PC (bottom) labelled for mGluR1α (18 nm immunogold), GABAB1 (red highlight, 6 nm) and CaV1.2 (blue highlight, 6 nm). Note the abundance of strong immunogold labelling under any condition.

| Electrophysiological  Property | Control  (18 cells,  14 mice) | Baclofen  pre-application  (14 cells,  8 mice) | P value |
| --- | --- | --- | --- |
| Membrane potential (mV) | -60.5 ± 8.8 | -59.1 ± 6.7 | 0.638 |
| Input resistance (MΩ) | 327.7 ± 118.6 | 218.2 ± 128.6 | **0.018 (*)** |
| Membrane time-constant (ms) | 29.9 ± 13.9 | 20.2 ± 12.8 | 0.051 |
| Capacitance (pF) | 93.7 ± 37.9 | 95.8 ± 42.4 | 0.884 |
| Rheobase (pA) | 40.0 ± 32.5 | 106.3 ± 69.2 | **0.003 (**)** |
| Voltage threshold (mV) | -41.6 ± 6.1 | -38.4 ± 6.5 | 0.162 |
| AP amplitude (mV) | 87.3 ± 16.6 | 74.8 ± 15.5 | **0.0378 (*)** |
| AP 20-80% rise-time (ms) | 0.15 ± 0.06 | 0.16 ± 0.07 | 0.865 |
| AP half-height width (ms) | 0.49 ± 0.17 | 0.47 ± 0.17 | 0.751 |
| AP maximum rise (mV.ms^-1^) | 370.3 ± 162.0 | 294.5 ± 124.9 | 0.158 |
| AP maximum decay (mV.ms^-1^) | 185.1 ± 84.2 | 169.7 ± 66.2 | 0.580 |
| Sag amplitude (mV) | 6.6 ± 4.1 | 4.4 ± 3.7 | 0.126 |
| Sag amplitude % of max | 19.0 ± 9.3 | 16.4 ± 13.0 | 0.514 |
| Adaptation index | 1.8 ± 0.8 | 1.4 ± 0.3 | 0.166 |
| FI Slope (AP. pA^-1^) | 0.19 ± 0.05 | 0.11 ± 0.06 | **0.007 (**)** |

*Table 1: Basal electrophysiological parameters of identified SST INs under control conditions and following baclofen pre-treatment*. Data are shown as mean ± SD for key electrophysiological parameters measured from hyper to depolarising current steps (-125 to +125 pA, 25 pA steps, 500 ms duration). Statistics shown as output from Student’s unpaired t-tests.

| Electrophysiological  Property | Control  (12 cells,  10 mice) | Baclofen  pre-application  (7 cells,  5 mice) | P value |
| --- | --- | --- | --- |
| Membrane potential (mV) | -64.4 ± 7.2 | -68.2 ± 8.5 | 0.312 |
| Input resistance (MΩ) | 106.4 ± 43.1 | 109.2 ± 34.2 | 0.887 |
| Membrane time-constant (ms) | 18.8 ± 4.7 | 10.3 ± 2.5 | **0.0004 (***)** |
| Capacitance (pF) | 199.2 ± 81.2 | 106.3 ± 51.4 | **0.015 (*)** |
| Rheobase (pA) | 200.0 ± 104.5 | 228.6 ± 95.1 | 0.561 |
| Voltage threshold (mV) | -40.4 ± 4.8 | -40.7 ± 4.0 | 0.878 |
| AP amplitude (mV) | 108.9 ± 11.9 | 111.1 ± 9.5 | 0.673 |
| AP 20-80% rise-time (ms) | 0.14 ± 0.03 | 0.14 ± 0.01 | 0.873 |
| AP half-height width (ms) | 0.86 ± 0.14 | 0.73 ± 0.05 | **0.032 (*)** |
| AP maximum rise (mV.ms^-1^) | 461.2 ± 117.4 | 426.2 ± 59.2 | 0.475 |
| AP maximum decay (mV.ms^-1^) | 92.6 ± 17.4 | 117.2 ± 8.6 | **0.003 (**)** |
| Sag amplitude (mV) | 6.0 ± 3.2 | 7.1 ± 3.8 | 0.491 |
| Sag amplitude % of max | 14.2 ± 4.9 | 15.9 ± 5.7 | 0.508 |
| Adaptation index | 4.65 ± 1.67 | 6.1 ± 3.6 | 0.262 |
| FI Slope (AP. pA^-1^) | 0.16 ± 0.05 | 0.14 ± 0.05 | 0.546 |

*Table 2: Basal electrophysiological parameters of CA1 PCs under control conditions and following baclofen pre-treatment*. Data are shown as mean ± SD for key electrophysiological parameters measured from hyper to depolarising current steps (-500 to +500 pA, 100 pA steps, 500 ms duration). Statistics shown as output from Student’s unpaired t-tests.

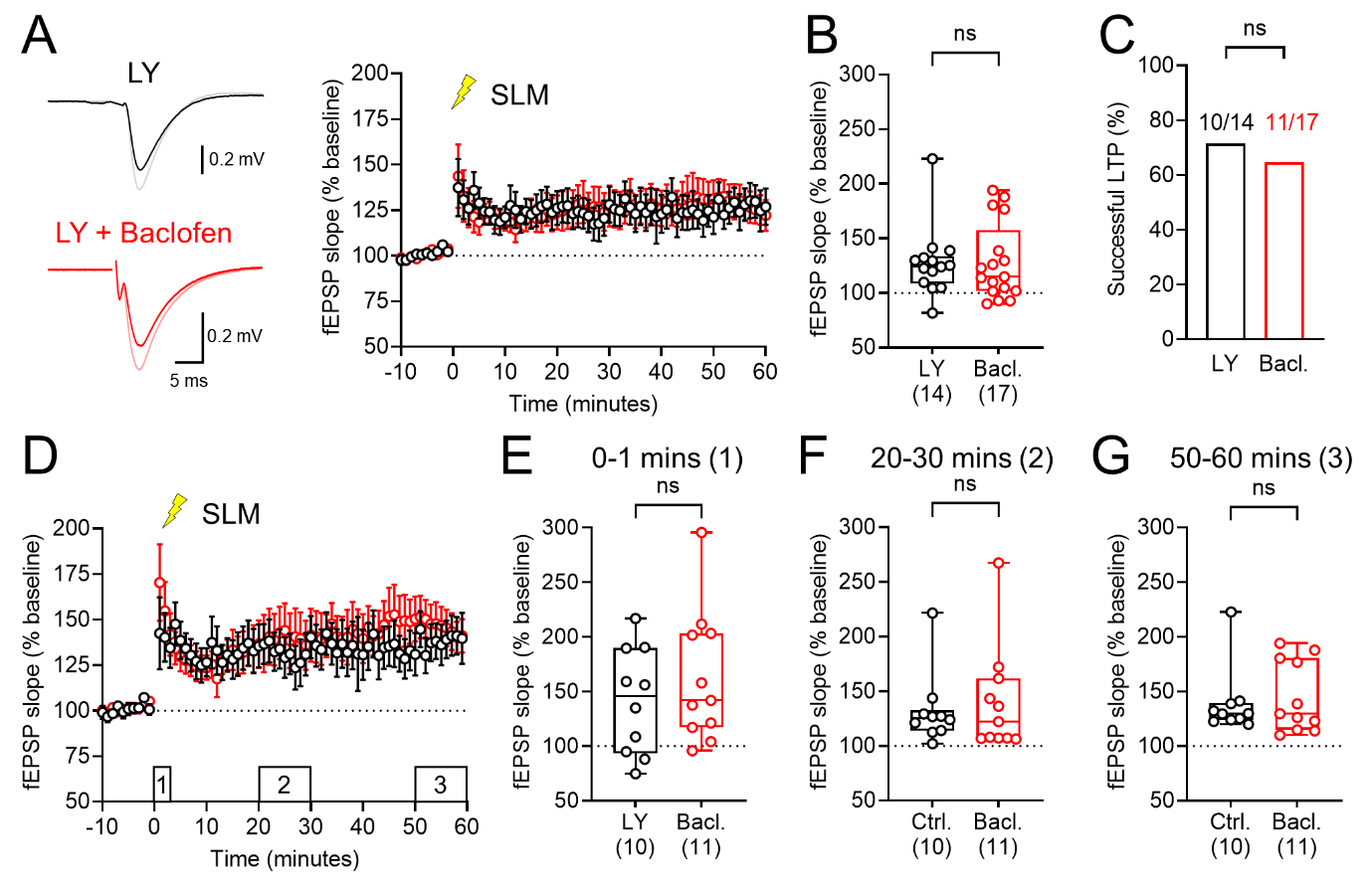
*Figure S4: Pharmacological inhibition of SST IN plasticity with mGluR1α antagonists prevents GABA_B_R internalisation induced facilitation of temporoammonic LTP*. A) Example fEPSP traces recorded in *str. L-M* in response to temporoammonic stimulation, and following induction of LTP in the *alveus* in the presence of LY-367,385 (LY) to antagonise mGluR1α in control (black) and 20 μM baclofen pre-treated slices (red)*.* Right, time-course of LTP in *str. L-M* in both slice conditions, recorded in the presence of LY. **B**) Quantification of the magnitude of LTP in all recordings performed in LY. **C**) Proportion of recordings in both control and baclofen pre-treatment that successfully induced LTP (>10% facilitation at 50-60 minutes post HFS) in the presence of LY. **D**) Time-course of successful LTP recordings induced in *str. L-M* in the presence of LY. Quantification of fEPSP facilitation in control and baclofen pre-treated slices at 0-1 (**E**), 20-30 (**F**), and 50-60 minutes (**G**) post *str. L-M* HFS, all in the presence of LY. Data is shown as either mean ± SEM (**A, D**), box-plots showing 25-75% box with median, with maximum range (**B, E, F, G**), pr proportions (**C**). Individual cell data is shown overlaid. Statistics shown as: ns – p>0.05, from Mann Whitney non-parametric tests or Chi-squared test (**C**).
